# Parallel Sensory Compensation following Independent Subterranean Colonization by Groundwater Salamanders (*Eurycea*)

**DOI:** 10.1101/2025.03.02.640989

**Authors:** Ruben U. Tovar, Brittany A. Dobbins, Nicholas R. Hartman, Sheena Leelani, Thomas J. Devitt, Dana M. García, Paul M. Gignac, David C. Cannatella, David M. Hillis

**Author notes:** Corresponding author: David M. Hillis, Department of Integrative Biology and Biodiversity Center, The University of Texas at Austin, Austin, TX 78712. **Competing Interest Statement:** The authors declare no competing interests. Classification: Biological Sciences: Evolution.

## Abstract

Lineages that have invaded subterranean environments have repeatedly evolved remarkable adaptations to life in darkness. However, observational and experimental studies in additional natural systems are needed to further our understanding of repeated evolution and convergence. In Texas, a radiation of groundwater salamanders (genus *Eurycea*), with independent invasions of subterranean karstic environments, offers an opportunity to investigate phenotypic convergence, parallel evolution, and the enhancement and regression of sensory systems. Adaptations to a troglobitic life in this clade includes morphological, behavioral, and physiological changes within and among species.

Intraspecific and interspecific variation in morphology to the selective pressures of life underground allows for detailed examination of physical, behavioral, and physiological changes associated with subterranean adaptation within a comparative phylogenetic framework.

We find a tradeoff between two sensory systems repeated across multiple subterranean *Eurycea* lineages: the degeneration of the eye and the expansion of the mechanosensory lateral line. The increase in anterior neuromast organs in subterranean lineages was positively correlated with the expression of paired box protein Pax-6, a conserved transcription factor important for vertebrate neurogenesis. Our results show a decreasing trend of PAX-6 labeling in the neuromasts of adult surface salamanders (*E. nana*) relative to the maintained labeling in subterranean species (*E. rathbuni*).

Our results suggest a tradeoff in resource allocation between the development of optic and anterior lateral line sensory systems in surface and subterranean salamander lineages and provide a starting point for future evolutionary developmental investigations examining the genetic underpinnings of adaptive, repeated evolution in a novel system.

**Significance Statement:** Under-explored subterranean environments in Central Texas harbor a phenotypically and taxonomically diverse radiation of groundwater salamanders, which we used to test the molecular and developmental bases for adaptive evolution in extreme environments. Using an integrative approach, we quantify the divergence and convergence of two sensory modalities: vision loss and mechanoreception. The divergent developmental and phenotypic tradeoffs have evolved several times in parallel among populations and species in this group. Understanding the evolutionary developmental processes responsible for these changes will further our understanding of adaptive, repeated evolution in a natural system.

## Introduction

Colonization of new environments can exert strong selective pressures on organisms, leading to new adaptive behaviors or traits (1). Subterranean habitats are considered extreme because they have little to no light and often limited food resources due to the lack of primary productivity (2). Organisms inhabiting these environments—termed stygofauna—are under common selective pressures that result in phenotypic convergence upon a suite of features characteristic of subterranean organisms, including reduced or absent eyes, reduced number of melanophores, elongation of appendages, and craniofacial modifications (3–6). In closely related lineages, shared phenotypes may result from parallel evolution in response to these common environmental pressures (7–10). Globally, stygofauna are represented by diverse metazoan taxa; among vertebrates, only teleost fish (13–16) and salamanders (4) have evolved stygobitic forms.

The karstic Edwards-Trinity aquifer system of west-central Texas, USA, is a vast network of subterranean aquifers comprised mainly of sandstone in the lower portions and porous limestone in the upper portions (11) with abundant surface springs. Weakly acidic precipitation facilitated the dissolution of Cretaceous limestone across the Balcones Fault Zone along the southern Edwards Plateau region. Since the Miocene, subterranean habitats have gradually formed from both geologic and hydrologic events. Climatic oscillations of alternating increased precipitation and drought (12) are associated with the evolution of an exceptionally diverse regional stygofauna.

The Edwards-Trinity groundwater ecosystem is home to 15 species of paedomorphic salamanders in the genus *Eurycea* (clade *Paedomolge*; 17). These salamanders are permanently aquatic and neotenic, retaining juvenile features into adulthood, including external gills and a broad tail fin. Some species are found in surface springs (e.g., *E. sosorum* and *E. nana*). One clade (the subgenus *Typhlomolge*) contains obligately subterranean species (*E. rathbuni*, *E. robusta*, *E. waterlooensis,* and an undescribed species). Other species, including *E. latitans*, *E. pterophila*, and *E. troglodytes*, inhabit both subterranean and surface spring habitats. Surface populations and species have fully pigmented skin and well-developed eyes. In contrast, obligately subterranean species exhibit features widely considered to be cave adaptations, such as a broad, flattened head, wide mouth, elongated limbs, greatly reduced numbers of melanophores, and vestigial eyes (6, 18–20). Species that have both surface and subterranean populations are highly variable in their subterranean-adapted traits (21); some populations lack melanophores and contain eyeless individuals but do not exhibit the extreme troglomorphism that characterizes *Typhlomolge* species, including dorsally-ventrally compressed head. Although not fully explored here, these polymorphic populations offer exceptional potential for identifying whether cave colonization is facilitated by genetic background, phenotypic plasticity, or both (22). Importantly, phylogenetic analyses support a history of repeated, independent colonization of subterranean habitats in *Paedomolge* (23–26; Fig. 1), which offers a compelling system for investigating the evolutionary developmental basis of adaptive evolution by quantifying the phenotypic changes that occur following colonization of an environment devoid of light.

**Figure 1.**
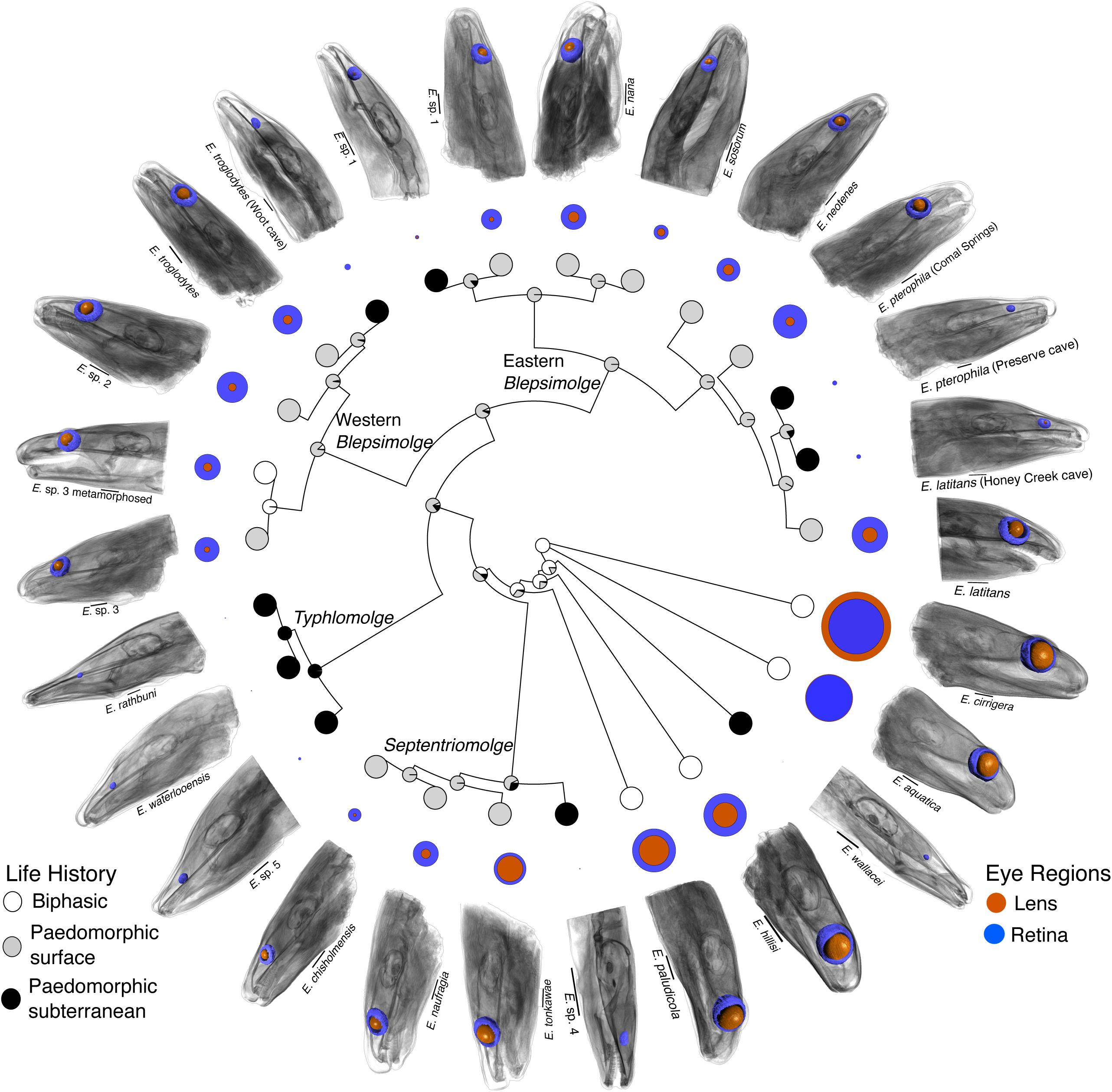
A phylogeny based on *cytochrome b* and *RAG 1* gene sequences. The reconstructed ancestral states were inferred from a one-rate model. Three life history states—biphasic (white circles), paedomorphic surface (gray circles), and paedomorphic subterranean (black circles)—were estimated and mapped onto each node. Six subterranean invasions are inferred within *Paedomolge*, and one outside this group (*E. wallacei*). The relative volumes of each eye region are plotted in nested circles around the perimeter of the phylogeny. The retinal volume (blue) is typically larger than the lens volume (orange) with the exception of *E. aquatica* and *E. cirrigera*, where the inverse is true, and in *E. aquatica* the lens is only marginally larger. The absence of a lens is observed in most subterranean species. DiceCT scans of exemplar heads follow the eye volume measurements and show the relative position and size in situ. Scale bars indicate 1 mm.

The degeneration or loss of eyes is one of the most salient features of cave-adapted organisms. Descriptions of subterranean salamander species have broadly categorized reduced eyes and pigmentation as convergent phenotypes (4). However, ocular and lateral line development have similar starting points among most vertebrates with early migration of placodes, including the lens and lateral line placodes, seen in tetrapods and fish (27). For example, eye formation in the eye of the Mexican blind tetra (*Astyanax mexicanus*) is developmentally conserved (5, 7) although it resides in complete darkness. Similar developmental conservation between the subterranean and surface *A. mexicanus* is observed, but eye developmental patterns diverge when cessation of growth and eventual apoptosis is initiated by the early-differential expression of Sonic Hedgehog (*shh*) compared to Paired box protein 6 (*pax6*) (7) in subterranean populations of *A. mexicanus*.

In salamanders, differing degrees of ocular regression are especially obvious in cave-adapted species of *Eurycea*, *Typhlotriton*, and *Proteus* (28–31). Although early developmental canalization of the vertebrate eye has been observed in several subterranean and fossorial species (7, 31, 32), the complex pathways by which functional eyes form include numerous opportunities for disruption. Thus, eye degeneration and loss may occur by different mechanisms among lineages (7, 32, 33). These mechanisms may leave the underdeveloped eyes in different stages of regression, but such a phenomenon has not been explored in a comparative framework in salamanders.

Without functional eyes, selection may enhance other sensory systems important in navigation, predator-prey interactions, and mate recognition as a compensatory mechanism (1, 34–36). The lateral line, a mechanosensory system present in fish and most larval amphibians, is likely to undergo compensatory evolution (37, 38). The lateral line system contains a series of epithelial mechanoreceptor organs (neuromasts; 39) found superficially (skin surface) in larval amphibians, or both superficially and in modified canals in fishes. Neuromasts respond to water pressure and movement (38). Inside each neuromast is a cluster of hair cells (*SI Appendix*, Fig. S1). From the apical portion of each hair cell projects tiered rows of stereocilia and a single kinocilium (*SI Appendix*, Fig. S2), extending into a jelly-like dome known as the cupula (40). Disturbances in the water create pressure waves that displace the cupula and bend the embedded cilia. The movement of the cilia results in changes in the membrane potential of the hair cell, which is communicated to attendant neurons and then to the brain. The lateral line system includes the anterior lateral line (ALL), which contains neuromasts of the head, and the posterior lateral line (PLL), comprising trunk and tail neuromasts. In the Mexican blind tetra (*Astyanax mexicanus*) there is variation in neuromast counts among populations; some subterranean populations have more ALL neuromasts than others, which in turn have more than their sighted, surface-dwelling ancestors (5, 7, 14, 36–38).

Aquatic salamanders have hundreds of neuromasts, making the lateral line system an important, non-visual sensory system (40, 41). Whether subterranean-adapted salamanders have expanded their lateral line in a similar fashion to cavefish has not been previously explored. To investigate whether eye degeneration and loss in *Eurycea* is correlated with expansion of the lateral line system, we quantified eye volume and the number and location of superficial neuromasts in developmental series of multiple surface and subterranean populations and species. We predict that subterranean salamanders will show parallel trends in both eye reduction and exhibit more neuromasts than surface species, with obligately subterranean species (*Typhlomolge*) showing the greatest degree of expansion.

## Results

### Phylogeny Estimation

Our phylogeny estimate is similar to others (42); a clade of paedomorphic species (*Paedomolge*) includes both surface and subterranean species, and some currently recognized species have populations with surface and subterranean ecotypes. Although we constructed an ultrametric tree, we did not time-calibrate it. However, comparison with figures in Stewart et al. 2024 (42) suggests that *Paedomolge* diverged from other *Eurycea* about 23 mya and subsequently diversified 19 mya.

### Evolutionary Patterns in Life History Traits (Ecotypes)

To quantify the rate of evolutionary shifts among the three ecotypes and estimate ancestral conditions on the phylogeny, we used ancestral reconstruction using four Mk models (43): one rate, six rate, ordered, and irreversible. We repeated these four analyses, adding a rate heterogeneity parameter to the models (see *SI Appendix*, Table S1 and Methods for details), and using a weighted average (44).

The transition from biphasic life history to paedomorphy occurred early during the divergence of the *Paedomolge* clade (Fig. 1). Within *Paedomolge,* at least six subterranean invasions occurred within the group, probably in the last 3 my, and at least one subterranean lineage is represented within each named subclade (Fig. 1). We infer this by adding stochastic character mapping (45, *SI Appendix*, Table S2) for the range of estimates of life history state transitions from the models we examined.

### Shape Differences and Homoplasy Among Ecotypes

Our analyses of continuous variation in shape variables included exploratory plots (*SI Appendix*, Figs. S4, S5, S6), assessment of phylogenetic signal, principal components (standard and phylogenetic) to explore overall shape differences among ecotypes, ANOVA and ANCOVA (standard and phylogenetic) tests of differences among ecotypes, and ancestral state reconstruction.

Phylogenetic signal in a trait is the tendency for related species to resemble each other more than they would if the trait were evolving at random. If the trait is evolving under Brownian motion, where phylogenetic signal should be high, then lambda equals 1. We assessed phylogenetic signal of raw morphometric variables and principal component scores using Pagel’s lambda (46) where 0 indicates no signal and 1 indicates strong signal; that is, the trait evolution tracks the phylogeny. Using likelihood-ratio tests, we tested the hypothesis that lambda = 1.0 and rejected it (P < 0.05) for the three eye variables, all three standard principal components, and two of the three phylogenetic principal components (*SI Appendix*, Table S3). We conclude that the size and shape of the retina and lens are not reliable indicators of phylogenetic relationships.

We performed both standard (non-phylogenetic) principal component analysis (PCA) and a phylogenetic principal components analysis (47) of the species means for Retina Total Volume, Lens Total Volume, and Snout-Gular length (a proxy for body size). The PCA shows that the first two axes explain 95% of the variance (*SI Appendix*, Fig. S7*A*). For all components of the PPCA, the interpretation of loadings and eigenvectors is almost identical to that of the PCA, so we discuss only the PCA (see *SI Appendix*, Table S4). PC 1 of the PCA is interpreted as a size factor because all loadings are positive (*SI Appendix*, Table S4). The ecotypes are easily separated along this axis (*SI Appendix*, Fig. S7*A*). PC 2 is a contrast between head size (SGL) and overall eye size (Lens Total Volume plus Retina Total Volume); species such as *E. waterlooensis* have large heads compared to their eye size (high score on PC 2). The subterranean species have low scores on PC 1 (overall small eyes) but span the entire range of PC 2 scores. *SI Appendix*, Fig. S7*C* shows substantial overlap among the ecotypes along this axis. PC 3, which explains only 4% of the variance, nonetheless separates the surface and subterranean species (*SI Appendix*, Fig. S7*B,C*); it is a contrast between Retina Total Volume and Lens Total Volume. Positive scores indicate a retina that is large relative to the lens and negative scores indicate the reverse. *SI Appendix*, Fig. S7*B* shows separation between the surface and subterranean ecotypes, but distribution of biphasic ecotype overlaps with both these two.

To quantify differences among ecotypes for each morphometric variable, we used generalized least squares (GLS) and phylogenetic generalized least squares analyses (PGLS) to conduct ANOVA and ANCOVA tests, for a total of four tests. We assessed ecotype differences in Retina Total Volume, Lens Total Volume, Eye Total Volume, PC 1, PC 2, and PC 3 and applied post-hoc comparisons (*SI Appendix*, Table’s S5-10). The interpretation generally did not differ whether SGL (head size) was used as a co-variate (ANCOVA) or not (ANOVA). The ecotypes were easily separated based on ocular measurements. Each of the six variables showed significant differences among the ecotypes under all four analyses, except in one case. For the three raw variables, the pairwise post-hoc comparisons were significant except for a few contrasts between surface and subterranean phenotypes (GLS), and surface and biphasic phenotypes (PGLS) in Lens Total Volume (*SI Appendix*, Table 7).

All PC 1 posthoc comparisons but one were significant (*SI Appendix*, Table 8). For PC 2, the GLS ANCOVA analysis was not significant, and posthoc comparisons for the other categories of analysis were mostly significant (*SI Appendix*, Table 9). For PC 3, none of the surface–biphasic comparisons were significant (*SI Appendix* Table S10), whereas most of the other contrasts were. Overall, the ecotypes can easily be distinguished by any of the morphometric variables, even though these phenotypes have evolved multiple times. This is easily seen, for example, in *SI Appendix*, Fig. S8 when the species are ordered by Eye Total Volume. The numerous overlapping branches indicate that the ecotypes, particularly the subterranean phenotype, do not sort out phylogenetically. To visualize evolution of the morphometric traits on the phylogeny, we mapped Eye Total Volume, Retina Total Volume, Lens Total Volume, and SGL onto the tree under a Brownian motion model (Fig. 2 and *SI Appendix*, Figs. S3, S9-11). These reconstructions emphasize the multiple reductions in eye size.

**Figure 2.**
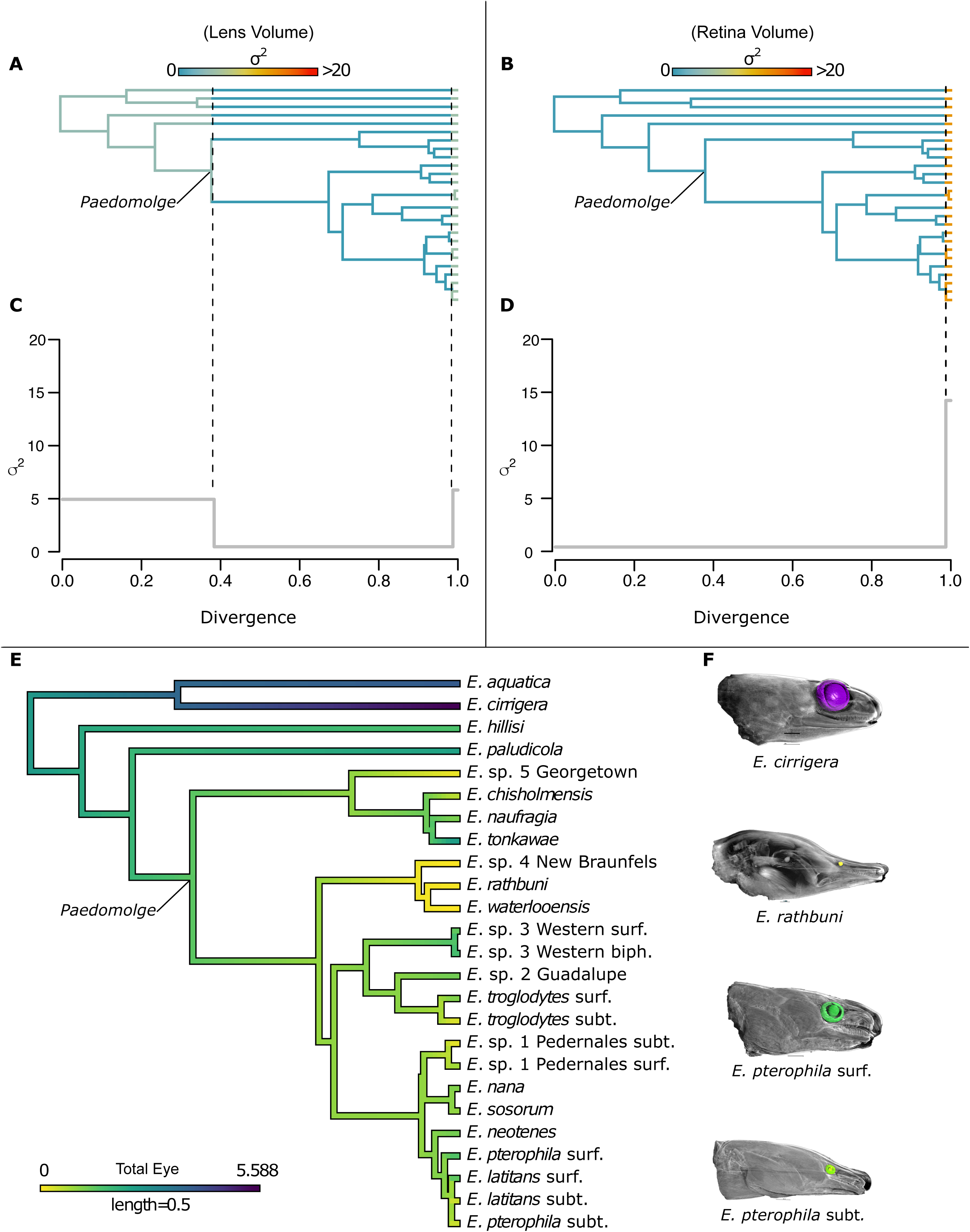
Trait rate modeling showing difference in regime models between the lens. (A) and the retina (B). A three-rate regime shift model for the lens (A) shows a decrease in divergence at the split between the *Paedomolge* clade, and an increase in divergence comparable to the initial rate (C) and corresponds to the terminal tips of the phylogeny (vertical dashed lines). A two-rate regime shift model (B) for the retina shows an increase in divergence (D) corresponding with the tips of the phylogeny (vertical dashed line). Ancestral state reconstruction of the Total Eye Volume under Brownian motion (E). Exemplar species with Total Eye Volume colored to reflect the species respective rate as observed by their respective tips (F).

### Morphological and Genetic Divergence Are Uncoupled

We compared the topology of the chronogram with that of a minimum evolution tree derived from a Euclidean distance matrix calculated from Retina Total Volume, Lens Total Volume, and Snout-Gular Length (*SI Appendix*, Fig. S3). The large number of crossing lines and the large normalized Robinson– Foulds distance (0.958) illustrates that the topologies are highly dissimilar. This indicates that shape evolution is largely independent of phylogeny.

### Phenotypic Evolution Undergoes Rate Shifts Across the Phylogeny

For Lens Total Volume, the three-rate model was best (AICw 0.601; Fig. 2). For Retina Total Volume, the two-rate model was best (AICw 0.794; Fig. 2). For Eye Total Volume, the two-rate model was best (AICw 0.513) but a combination of the two– and three-rate (AICw = 0.365) models accounted for 0.878 of the weight *SI Appendix*, Table S11) (*SI Appendix*, Fig. S12-15). It is notable that the two-rate models of Eye Total Volume and Lens Total Volume show a drastic increase in σ^2^ very near the tips of the tree. When a third rate is added to the model, there is a distinct decrease in σ^2^ at the level of the ancestor of *Paedomolge* (Fig. 2*A*). The rate shift plot of PPCI (*SI Appendix*, Fig. S15) provides a summary of the rate shifts of the eye variables. The results indicate that the factors contributing to shifts in the evolutionary rates of these traits are not specific to lineages. Specifically, the dramatic increase in σ^2^ at the tips of the tree suggests recent changes in multiple lineages.

### Eyes of Subterranean Species Undergo Reduction Early in Development

The structure of the retina has not been described in most of the species. The retina of surface ecotypes is composed of seven layers we identify as the retinal ganglion cell layer, inner plexiform layer, inner nuclear layer, outer plexiform layer, outer nuclear layer, photoreceptors, and pigment epithelium (*SI Appendix*, Fig. S16) (6). We identified the composite of all layers as the retina for all downstream segmentation and volume measurements of adult specimens. We also identified the ciliary body as an extension from the distal portions of the retina, attaching to the lens in adults. We assume that the ciliary body functions similarly in *Paedomolge* relative to other vertebrates, including other salamanders. In other salamanders, the ciliary body contributes to mechanically displacing the lens for accommodation. The diffusible iodine-based contrast-enhanced computed tomography (diceCT; 107) scans of early salamander development do not clearly distinguish the boundary between the posterior ciliary body and anterior retina. Therefore, we included the developing ciliary body, known as the ciliary marginal zone (48) in the retinal measurements of all early developmental stages. The lens was identified as a large spheroid cupped by the retina, and in the adults, is attached to the ciliary body. An optic nerve can be identified in all the adult and developmental specimens. It appears as a thin, relatively well-stained nerve bundle emanating from the retina.

To trace the time course of eye reduction during development, we measured the volumes of the lenses and retinae of larvae (a minimum of three individuals per stage) 1-, 2-, 3-, 4-, and 5-months post-oviposition (mpo), as well as in adults (adults described above), and compared the volumes among three surface taxa (*E. nana*, *E. pterophila*–Comal Springs, and *E. sosorum*) and three subterranean taxa (*E. rathbuni*, *E. latitans*–Honey Creek Cave, and *E. pterophila*–Preserve Cave) using a generalized linear model (*SI Appendix*, Table S12). An analysis of variance (ANOVA) was conducted on the six respective species at each stage of development (*SI Appendix*, Table’s S13-18*A*-*D*; Fig. 3*B*). At 2 mpo, differences emerged between ecotypes and became more striking as development continued, as illustrated in the Tukey’s post hoc distribution bars and estimated marginal means (Fig. 3*A*, *SI Appendix*, Fig. S17). From stages 2 mpo to adult, Eye Total Volume varied between phenotypes, with increasingly greater eye volume in surface taxa relative to subterranean taxa (Fig. 3*A*&*B*).

**Figure 3.**
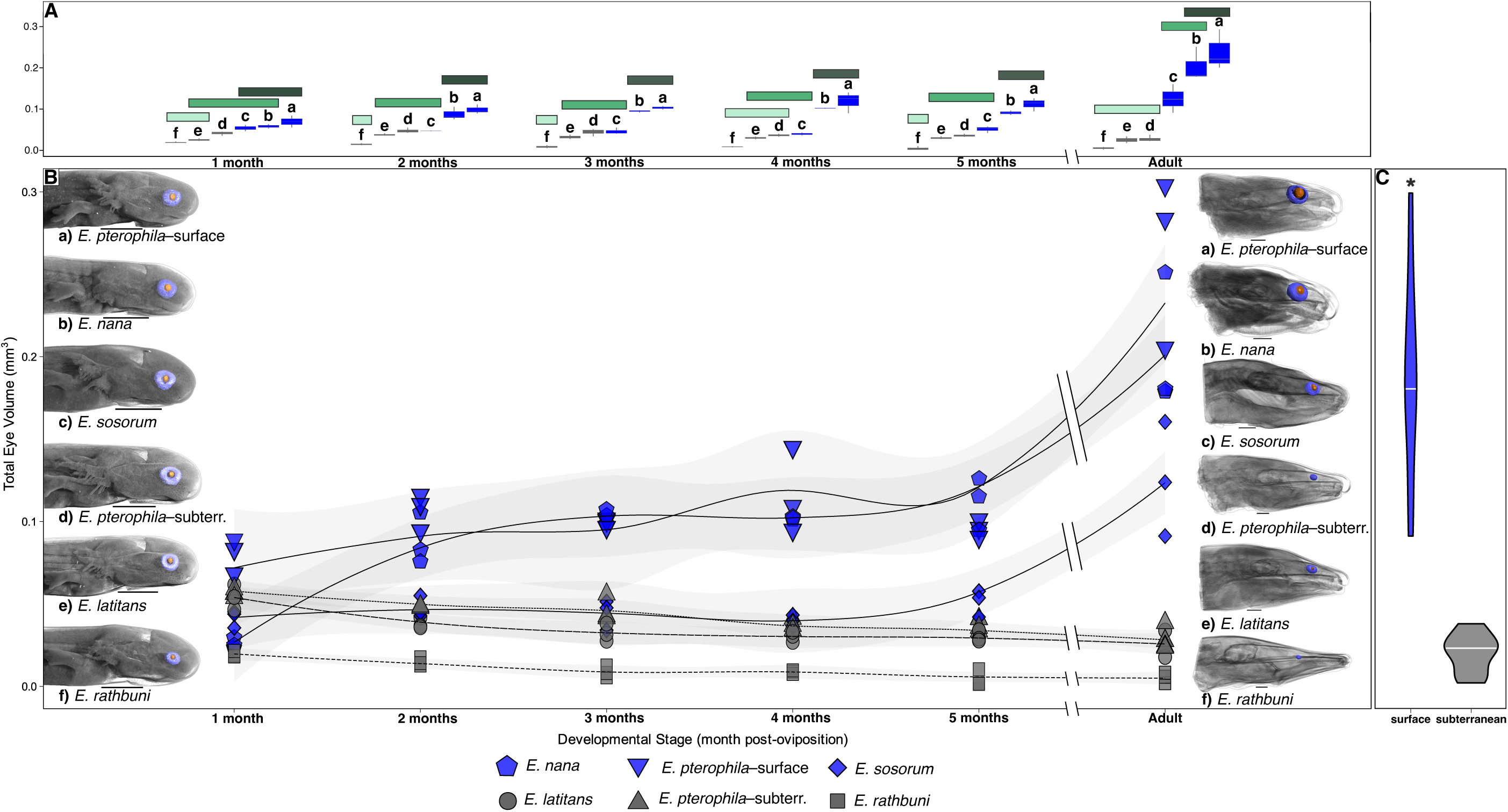
(A) Generalized linear regression was used to measure and plot the developmental trajectories of eye volume through a developmental series of six taxa of *Eurycea* withTukey’s grouping in three shades of green, and box plots associated with surface (blue) and subterranean (grey) species. Taxa include three surface populations (a: *E. pterophila*– surface; Comal Springs; b: *E. nana*; c: *E. sosorum*) and three subterranean populations (d: *E. pterophila*– subterranean; Preserve Cave; *E. latitans*–Honey Creek Cave; f: *E. rathbuni*). Eye volumes are plotted for each developmental stage (1-5 months post-oviposition and adults; n=3 for each stage and taxon). (B) Standardized adult eye volumes between surface and subterranean species are significantly different (the retina in blue and lens in orange). Scale bars indicate 1 mm. Already at one month, distinctions are starting to be made between different ecotypes these differences become more striking as development continues. Results of a generalized linear regression with adult only eye volume are show in a violin plots in panel (C) with significant difference between surface (blue) and subterranean (grey) species.

There was a significant difference in volumes of the adult retina and lens among six species (*E. rathbuni*, *E. latitans*, *E. pterophila*–PC, *E. pterophila*–CS, *E. nana*, and *E. sosorum*) based on diceCT images (*SI Appendix*, Table. 16*A*-*D*). Total eye volume is significantly different among species (*P* < 3.86e-07, *SI Appendix*, Table. 16*A*), and between ecotypes (Fig. 3*C*, post-hoc Tukey’s test; *SI Appendix*, Table. 16*C-D*), but not among species within ecotypes. The three subterranean species have less lens and retinal tissue than surface species. Among the subterranean species, the lens tissue volume is reduced or in most cases absent in adults (Fig. 1; blue versus orange circles). This is different from metamorphosed individuals outside the *Paedomolge* clade. In *E. aquatica* and *E. cirrigera* the adult lens volume is greater than the retinal volume.

All subterranean adult individuals had reduced retinae and adult *Eurycea rathbuni*, *E*. sp. 5, *E. waterlooensis*, and *E. wallacei* had no lenses (Fig. 1). *Eurycea latitans* and *E. pterophila* (PC) are from populations of polymorphic species (i.e., populations with multiple discrete phenotypes) and have greater variation in lens development; in these populations, some individuals retained underdeveloped lenses and retinae.

### PAX6 Localizes to Eyes and Neuromasts

To determine whether eye reduction in subterranean salamanders was related to diminished expression of the gene encoding the developmental transcription factor Paired-box 6 (PAX6), recognized as a master regulator of eye development, we compared immunolabeling for PAX6 between *Eurycea rathbuni* and *E. nana* at 1– and 3-mpo and in adults. We examined at least 3 individuals for each species at each stage of development using a polyclonal anti-PAX6 antibody, validated in mammalian species and recently in *E. latitans* (94). Western blot analysis indicated that this antibody labeled a band at the appropriate molecular weight in tissue homogenates obtained from *E. nana* and *E. rathbuni* embryos stages 37–41 (*SI Appendix*, Fig. S18). We observed no statistically significant difference in PAX6 labeling in the eye between *E. nana* and *E. rathbuni* through the three early stages of development (Fig. 5). However, there is a significant difference (*P* < 0.05) in the PAX6 labeling of neuromasts between adult *E. rathbuni* and *E. nana* (Fig. 5). Specifically, *E. nana* adults showed lower levels of PAX6 labeling compared to *E. rathbuni* (Fig. 5*B*, 5*G*). Moreover, we observe a negative relationship between PAX6 labeling through *E. nana*’s development relative to *E. rathbuni*. Localization of this transcription factor was both nuclear and extranuclear with particularly intense labeling in the apical appendages of the hair cells (Fig. 5*B*–*F*).

**Figure 4.**
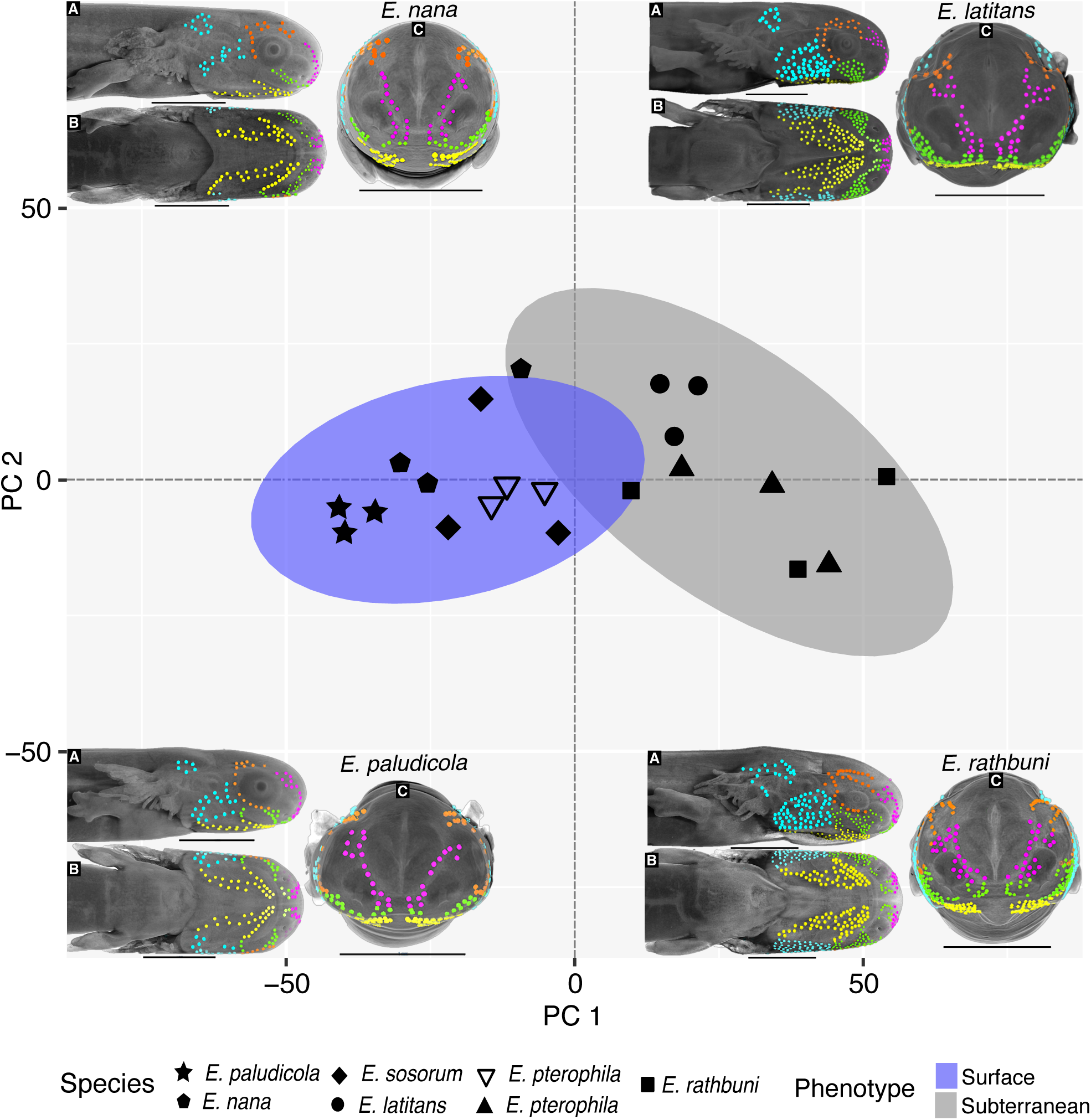
Principal components analysis of neuromast counts among seven species of *Eurycea* at 1-month post-oviposition. PC 1 (positive loadings for all neuromast counts) accounts for 82.4% of the variance and indicates higher neuromast counts for subterranean compared to surface species). PC 2 accounts for 11.2% of the variance and is largely a contrast between maxillary and mandibular neuromast counts (*SI Appendix*, Fig. S4). Exemplar species and their respective neuromasts are illustrated for each quadrant of the PCA (clockwise: *E. nana*, *E. latitans*, *E. rathbuni*, and *E. paludicola*). Sagittal (A), ventral (B), and frontal (C) views are shown for each exemplar species. Neuromasts are shown in situ, in postorbital (light blue), orbital (orange), mandibular (yellow), maxillary (green), and nasal (purple) regions. Scale bars indicate 1 mm.

**Figure 5.**
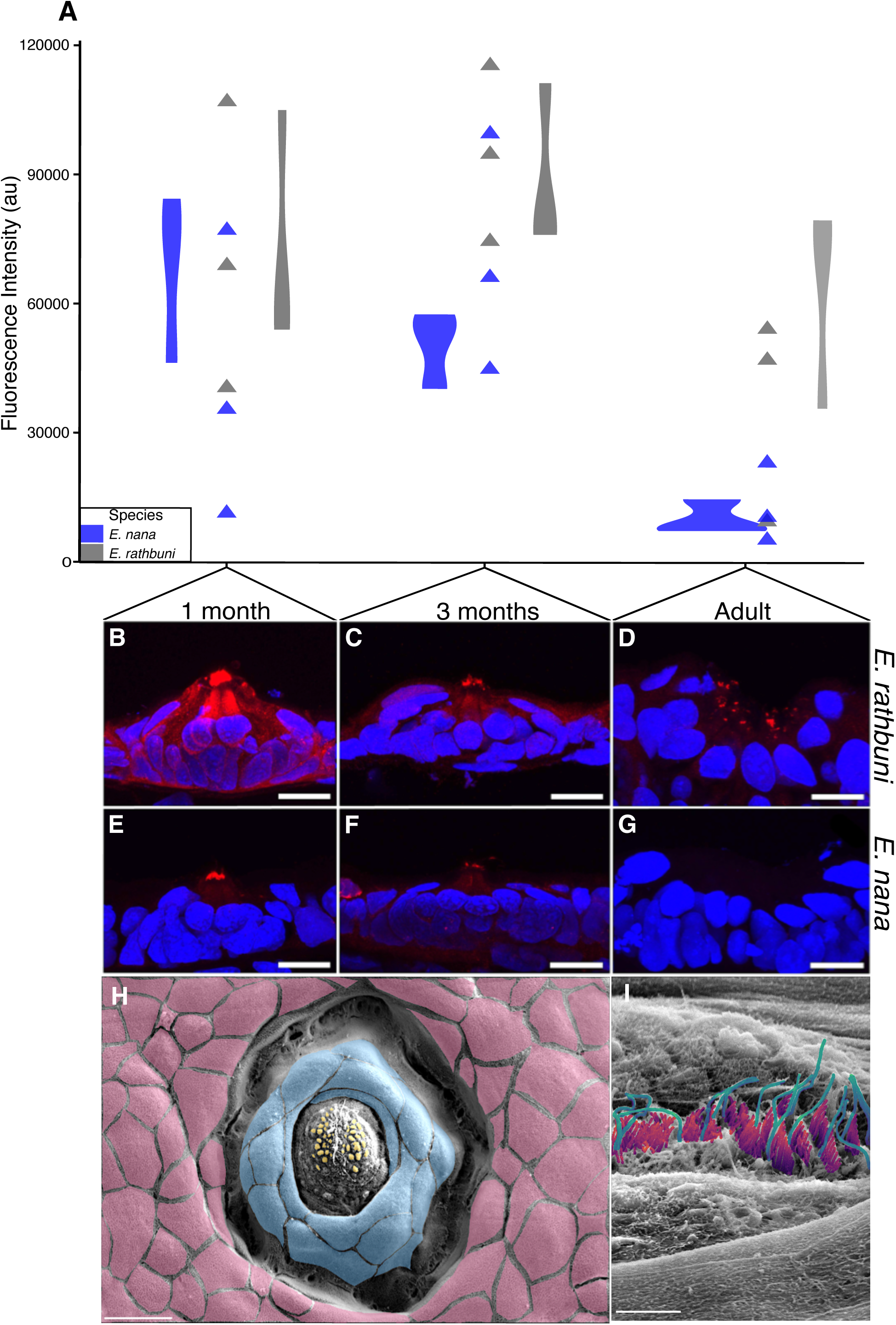
Differential PAX6 labeling in eyes (triangles) and neuromasts (violin plots) through ontogeny. (A) ANOVA points distributions show three developmental stages between two species (*E. rathbuni* in gray and *E. nana* in blue). (B–G) Sections of neuromasts labeled with DAPI nuclear stain (blue), and anti-PAX6 (red), white scale bars indicate 1 µm. An SEM image of *E. rathbuni* neuromast microstructure (H) identifying epethilial cells (pink), mantle cells (blue), and support cells (orange), scale bar equals 20 µm. A higher magnification of the same neuromast (I) shows kinocilia (aqua blue), and stereocilia (magenta), scale bar equals 5 µm.

### Subterranean Species Have More Anterior Neuromasts

The observation of differences in PAX6 labeling in neuromasts of *Eurycea rathbuni* compared to *E. nana* led us to ask whether the lateral line system of subterranean salamanders was enhanced relative to surface ecotypes.

Neuromasts appear as small, round or elliptical indentions on the skin’s surface in the diceCT renderings, matching Lannoo’s description (49). To corroborate our identification of neuromasts in 1-month post-oviposition specimens, we produced a higher resolution diceCT scan (*SI Appendix*, Fig. S1*B*, S1*C*) and images using scanning electron microscope of an adult *Eurycea rathbuni* (Fig. 5*H*), enabling us to visualize superficial and subcutaneous structures indicative of neuromasts (50). Mapping and quantifying neuromasts of the anterior lateral line of salamanders from seven species of *Eurycea* (four surface and three subterranean species; Fig. 4) revealed a significant difference in the number of ALL neuromasts between surface and subterranean salamanders (one-way ANOVA; F(_6,14_) = 16.10, *P* < 0.001). There are significantly fewer ALL neuromasts in the four surface salamander populations than in two of the three subterranean populations (*P* < 0.05). The remaining subterranean representative (*E. latitans* HC) did not have a significantly different number of neuromasts than surface *E. pterophila* CS individuals (*P* = 0.081).

Principal components analysis (PCA) of regional neuromast counts largely separated surface and subterranean species along PC 1, which accounted for 82.4% of the variation and had positive loadings for all neuromast counts (Fig. 4). PC 2 accounted for 11.2% of the variation and separated samples by mandibular and maxillary neuromast counts, with the greatest differences among the subterranean species (Fig. 4; *SI Appendix*, Fig. S19). We further examined neuromast structure with scanning electron microscopy, revealing superficial structures of the neuromast, including kinocilia and stereocilia in both *E. nana* and *E. rathbuni* (*SI Appendix*, Fig. S2). Additional structures are also evident in the SEM images, for example, epidermal microvilli on the surface of *E. nana*’s skin and a pit organ in *E. rathbuni*. Histological examination of tissue sections revealed several hallmark neuromast structures, including hair cells, mantle cells, and cilia (*SI Appendix*, Fig. S1 & S2).

## Discussion

### Repeated Eye Reduction and Cave Colonization

Our phylogenetic analysis (Fig. 1) indicates that the transition from a biphasic life history to paedomorphy occurred early in the *Paedomolge* clade, with one reversal. Invasions of subterranean environments, and the evolution of the subterranean phenotype (eye and pigment reduction or loss), have occurred independently at least six times, mostly within the past few million years (Fig. 1). There have been multiple convergences and reversals in ecotypes (estimated using stochastic mapping, under any of the models).

Within a phylogenetic framework, the three ecotypes (biphasic, paedomorphic surface, and paedomorphic subterranean) are readily distinguished using the morphometric variables (either raw measures or PCs), using phylogenetically informed ANOVA and ANCOVA. Adding SGL as a covariate makes no difference, suggesting that SGL is not a good estimator of overall size. None of the continuous traits we measured were indicators of phylogenetic relationship, supporting the conclusion that eye reduction and lateral line augmentation arose independently in multiple lineages. This is evidenced by the phylogenetic signal analysis and the co-phylogenetic analysis of morphology and genetic distance. The rate-shift analyses suggest that the some of the factors contributing to shifts in the evolutionary rates of these traits are not specific to independent lineages. Specifically, the dramatic increase in σ σ^2^ at the tips of the tree suggests an external influence, such as the recent repeated invasion of cave habitats.

### Lateral Line Augmentation

In addition to eye reduction or loss, we found repeated sensory augmentation across the multiple lineages of salamanders that have independently colonized subterranean habitats. This augmentation is manifested as increased numbers of neuromasts in the anterior lateral line (ALL) system in subterranean ecotypes. We interpret the co-occurrence of eye loss and ALL enhancement as a mechanism of sensory compensation in which a reduction of a dominant sensory system led to the enhancement of another sensory system, thereby ensuring individuals retain an ability to interpret their environment.

For compensation to have occurred, the sensory modality dominant in the ancestral state must have first been diminished. Augmentation of a previously non-primary sensory modality then occurs. This process is likely to occur following a major ecological transition, in which a new environment favors one or more secondary sensory modalities (51, 52, 53). We propose that this sequence most parsimoniously describes the phenotypic transition exemplified by cave-dwelling salamanders. Namely, vision is important in feeding, navigating, and finding mates in surface salamanders (54), which is suggested by the photic environment and ancestral presence of well-developed eyes for *Eurycea*.

Reliance on light becomes impossible underground, irrespective of whether eye loss occurs immediately upon habitat transition. This reduced ability to use vision to interact with the environment may facilitate an alternative sensory system to dominate. For aquatic animals, it is common to evolve or augment mechanosensory systems capable of sensing (55, 56), interpreting (57, 58), and mapping changes in water pressure (59). Such augmentation is exemplified by distantly related groups with convergent mechanoreceptor systems such as the neuromasts of fishes (36, 37), the dome pressure receptors of crocodilians (60), and the vibrissae of aquatic mammals (61).

We found that numbers of neuromasts increased on the head, mandibular, maxillary, post-orbital, and nasal regions (Fig. 4; Fig. S19). The increase in postorbital neuromasts may provide enhanced sensory capability by extending coverage of the posterior region of the head, especially considering the expanded surface area offered by dorsoventrally compressed and lateral expansion of the heads of most subterranean species (19). An increase in neuromasts posterior to the eye may assist in predator avoidance or mating. Like the mandibular, maxillary, and post-orbital regions, the nasal and orbital regions show increased neuromast counts in subterranean species, but to a lesser degree. Maxillary and mandibular concentrations presumably facilitate prey detection in an austere subterranean environment. The vibrations released by potential prey items (isopods, amphipods, and other invertebrates) are signals readily interpreted by neuromasts (95). Chemosensory organs could also be important in this regard.

We show that a pronounced reduction in ocular development correlates with increased neuromast counts in subterranean salamander lineages. Thus, the augmentation in neuromast numbers among subterranean salamanders provides an enhanced sensory modality compared to the ancestral condition. Sensory compensation could be accomplished through paedomorphy. Mid-Miocene climatic warming and associated aridification in Texas presumably favored a transition to paedomorphy in the ancestor of *Paedomolge* (12, 26). The descendant species in this group are obligately aquatic and retain larval characteristics (e.g., gills and tail fin) adaptive for an aquatic life history, except in rare instances in one population of one species (*E. troglodytes*).

### Role of Heterochrony

Heterochrony—the displacement of a trait’s development in time relative to its ancestral state—likely played an important role in trait transitions from surface to subterranean phenotypes. However, the molecular mechanisms that produce paedomorphic and peramorphic (i.e., overdevelopment of a trait in derived adults; 62, 63) traits are poorly understood. Furthermore, paedomorphy could be manifested to different degrees. For example, perhaps the failure of eye development reflects an extreme form of paedomorphy in which heterochronic maturation of the gonads still occurs while the rest of the body is left in a juvenile state or, in the case of the eye, possibly an embryonic state.

Similarly, paedomorphy may also play a role in enhanced lateral line development. Although the lateral line system is lost in most tetrapods, it is present in larval amphibians and paedomorphic salamander species (e.g. *Ambystoma mexicanum*; 64). The distribution and arrangement of the lateral line changes after metamorphosis, and neuromasts are lost altogether in some direct-developing salamander species (e.g. *Desmognathus aeneus* and *D. wright*; 65). However, the harbingers of neuromast development—afferent lateral line neurons—are still present in these *Desmognathus* species, suggesting a degree of neuro-anatomical paedomorphy (66). Interestingly, facultative metamorphosis occurs in at least two species within *Paedomolge* (*E. troglodytes* and *E*. sp. 3), and our observation of facultative metamorphosis in *E*. sp. 3 is shown in Fig. 1.

Surface paedomorphic salamanders have a suite of characteristics that are often associated with surface-dwelling organisms, including pigmentation, relatively small and robust limbs, and fully developed eyes (6, 23, 25). We observed lenses, well-developed retinae, and optic nerves in all surface-dwelling species. Furthermore, the lens and retina of surface species are markedly larger than those observed in subterranean *Eurycea* (Fig. 1). Most obligately subterranean *Eurycea*, such as *E. rathbuni* and *E. wallacei* (67), have drastically reduced eyes (Fig. 1), a characteristic that reflects subterranean-adapted morphology and which is observed in other subterranean salamanders (e.g., *E. spelaea* and *Proteus anguinus*; 30, 31, 68, 69), subterranean fish (e.g., *Astyanax mexicanus*, *Phreatichthys andruzzii*, *Sinocyclocheilus anophthalmus,* and *Pimelodella kronei*; 7, 70-72), fossorial organisms (e.g. moles and snakes; 33, 64), as well as invertebrates (74, 75). Furthermore, a number of *Eurycea* species (*E. rathbuni*, *E*. sp. 5, E. sp. 4, *E. waterlooensis*, *E. troglodytes*, and *E. wallacei*) exhibit a few vestigial retinal layers surrounded by pigment epithelium (Fig. 1; 6, 29, 41, 76). Pigmentation surrounding the eye suggests light would be unable to pass through choroidal pigment or pigment epithelium to be absorbed by any existing photoreceptors, a finding suggested for *E. rathbuni* (6) and *P. anguinus* (77). Interestingly, the optic nerve is retained in both species, suggesting a possible sensory, but not necessarily image-forming, function (66).

### Role of *Pax6*

Paired box 6 gene (*Pax6*) controls eye development across vertebrates (33, 78–80), as does its homolog (*eyeless*) in invertebrates (65, 81, 82). The product of this gene, PAX6, is a well-known transcription factor and is often observed as an endonucleic protein that plays a critical spatiotemporal role throughout development (83–85). Interestingly, PAX6 has also been documented in tissues other than eye, transiently through development (86, 87, 88), in cell culture (89), and during regeneration (90, 91). Extensive work by Jeffery (5) compared eye development between surface and subterranean phenotypes of *Astyanax mexicanus*; he showed that the overexpression of *shh* down the midline of the subterranean tetra leads to the down-regulation of *Pax6* (7). This down-regulation results in apoptosis and underdeveloped eyes in subterranean *A. mexicanus*, and a similar trend has been noted in the eye of *E. rathbuni* (6), although the paucity of specimens precluded quantification in that study. Here, we add additional, later developmental stages. We found labeling in the developing eye tissue of *E. rathbuni* is not significantly different from that of *E. nana* through ontogeny (*SI Appendix*, Fig. S6). This finding aligns more closely with observations in another cavefish species (*Phreatichthys andruzzii*), in which the expression of *Pax6* during early development does not deviate from canonical expression, suggesting that an alternative molecular target, and not primarily PAX6, is responsible for eye reduction (92, 93).

In contrast to our finding of that PAX6 labeling in eyes did not statistically significantly differ among stages. Through development of both a surface and a subterranean salamander, we found its abundance waned in the neuromasts of *Eurycea nana,* resulting in a negative trend through the three stages. We recently reported our observation of PAX6-labeling in the neuromast of *E. rathbuni* and *E. nana* (94), and here expand those observations by quantifying the labeling of PAX6 in the neuromasts of the subterranean *E. rathbuni* and surface *E. nana* through three developmental stages. Whether the persistence of PAX6 labeling in *E. rathbuni* neuromasts is a result of their augmentation requires further testing; however, an increase in the number of ALL neuromasts was evident in all the subterranean forms (*E. rathbuni*, *E. latitans–*HC, and *E. pterophila–*PC) we examined. Likewise, the subterranean populations of dimorphic species (*E. latitans* and *E. pterophila*) have more neuromasts (Fig. 4), as do some subterranean populations of *Astyanax mexicanus* (7).

## Materials and Methods

### Sample Collection and Sequencing

Salamanders were either collected from the wild or donated by the San Marcos Aquatic Resources Center (SMARC), USFWS, San Marcos, Texas (*E. nana, E. pterophila*–Comal Springs, *E. rathbuni,* and *E. sosorum*) and the City of Austin (*E. sosorum and E. waterlooensis*). We assembled a dataset of 26 cytochrome b (cytb) sequences and 17 recombination activating gene 1 (RAG1) from published GenBank sequences (n=24) and unpublished sequences that we generated (n=2; Fig. 1; *SI Appendix*, Table S19). We isolated DNA using spin column kits (Qiagen) and amplified a fragment of cytb (17). Sanger sequencing was performed by Eton Bioscience, Inc. using the same primers that were used in PCR. Geneious Prime 2024.0.3 (GraphPad Software, LLC) was used to assemble forward and reverse sequences and aligned using MAFFT 4.475 (96).; bases with more than a 1% chance of an error were trimmed from the sequence ends (*SI Appendix*, Table S19).

### Phylogenetic and Comparative Analyses

We performed a maximum likelihood phylogenetic analysis using IQ-TREE 2.3.4 (97) with ten replicates under the edge-proportional partition model (98) with 100000 ultrafast bootstrap replicates (99) to assess support (*SI Appendix*, Tables S9-10). Each analysis was repeated 10 times to ensure consistency. All currently recognized species of *Paedomolge* were included. The phylogram was converted to a chronogram with a tip to root distance of 1.0 using the R function ape::chronos().

Because some of the species were polymorphic for ecotype (particularly, species with some surface populations and subterranean populations), we included one representative of each ecotype in the phylogenetic analysis, even though there may be multiple ecotypes within a species. This yields a conservative estimate of the number of state changes.

Ancestral state reconstruction on both continuous and discrete traits was performed using the R packages *ape version 5.8* (100) and *phytools* version 2.3-0 (101). For details of the models, see the *SI Appendix*.

### Stochastic Character Mapping

To quantify the frequency of character-state changes, stochastic character mapping (45), a Markov Chain Monte Carlo procedure that simulates a phylogenetic map of history of a discrete character under a specified model, was done using phytools::make.simmap(). We analyzed the ecotype data under four Mk models and four corresponding Mk gamma models; the latter include a rate heterogeneity parameter. We excluded *E. wallacei* because other *Eurycea* species in this region of the phylogeny are poorly sampled, and inclusion of extreme values of *E. wallacei* seemed to bias the trait estimates. Additional details are given in the *SI*.

### Shape Differences Among Ecotypes

Principal component analyses were performed on the species means of the untransformed variables Retina Total Volume, Lens Total Volume, and Snout-Gular Length. Standard (PCA, nonphylogenetic) analyses were done using the stats::prcomp() with a correlation matrix. Phylogenetic principal component analyses (PPCA) were performed using phytools::phyl.pca(). Results are in *SI Appendix*, Fig. S7.

The nlme::gls() function was used to perform generalized least squares (GLS) and phylogenetic generalized least squares analyses (PGLS) under ANOVA and ANCOVA (SGL as the covariate) models, for a total of four tests for each variable. Life History was the predictor. The ape::corbrownian() function was used to specify within-group phylogenetic correlation structure for the PGLS analyses. We assessed ecotype (life history) differences in Retina Volume, Lens Volume, Eye Total Volume, PC 1, PC 2, and PC 3) and applied post-hoc comparisons using multicom::glht() and the “Tukey” option (*SI Appendix*, Tables S5-10).

### Phylogenetic Signal

The amount of phylogenetic signal (lambda) in the continuous traits was estimated using phytools::phylosig() (46, 102). The continuous variables were mapped onto the chronogram under the Brownian motion model using phytools::contMap(). Figure 2E shows the map for Eye Volume; the maps for other continuous variables are in (*SI Appendix*, Fig. S9-11).

### Chronogram

Raw values (mm^3^) for Eye Total volume by species (y-axis) were plotted against scaled genetic divergence (x-axis) and superimposed on the chronogram, using phytools::phenogram(). Symbols are colored by life history (white = Biphasic, gray = Surface, black = Subterranean).

### Rate-Shift Analyses

The rate-shift analyses (102) estimated points in time on the chronogram where the rate of character change, as measured by σ ^2^ (evolutionary rate of under Brownian motion) has shifted across all lineages simultaneously rather than along one branch. The algorithm identifies optima for one, two, three etc. points, at which there is a shift in the value of σ ^2^. These shift points delimit so-called regimes, the slices of time during which the σ ^2^ is constant. We analyzed each variable for four regimes and used AIC weight to determine the best fit model.

### Cophylogenetic Analysis of Morphology and Genetic Divergence

This analysis compares two trees, the DNA chronogram used for other analyses, and a phenotypic Minimum Evolution tree (ape::fastme.bal(); (103) calculated from a standardized Euclidean distance matrix (factoextra::get_dist(). Three variables were used: Retina Total Volume, Lens Total Volume, and Snout-Gular Length (SGL). The normalized Robinson-Foulds distance, which describes similarity of the topologies, was calculated using phangorn::RF.dist(). The distance ranges from 0 to 1; the calculated distance (0.958) means that the topologies are very different. The diagram was produced by the phytools::cophylo() function.

### Specimens

#### Early Development

All animal manipulations were approved by The University of Texas at Austin Institutional Animal Care and Use Committee (IACUC) under AUP-2021-00090. A developmental series was collected for four species, representing one subterranean species (*Eurycea rathbuni*) and three surface species, *E. nana*, *E. sosorum*, and *E. pterophila* from Comal Springs. Three individuals representing the above species for each stage were collected from the San Marcos Aquatic Resource Center (SMARC), Texas, United States Fish and Wildlife Service (USFWS) and incubated in an environmental chamber at 21.1°C until they were euthanized using Salamanders were euthanized using 0.2% MS222. A developmental series was obtained for two additional species and one population representing the subterranean phenotype of *E. pterophila* from Preserve Cave.

These three species were collected from in-house breeding populations at The University of Texas at Austin, maintained, and euthanized as described above. Three individuals for each species or population at each stage were collected. In total, a developmental series for six lineages of Texas *Euyrcea* were collected including both forms (surface and subterranean) of a polymorphic population of *E. pterophila* resulting in 108 specimens. Staging was determined by morphology (104), and embryos were maintained at constant temperature with no lighting except when fed by hand once yolk was fully absorbed after two months post oviposition. Light exposure during feeding was estimated to be approximately 10–15 minutes, twice a week. All specimens were collected and transported under Texas Parks and Wildlife scientific permit number SPR-0119-004, and State Park permit number 57-22.

#### Adult Specimens

Adult salamanders of fourteen species were collected including *E. rathbuni* (n=3), *E. nana* (3), *E. sosorum* 3), *E. pterophila* from Preserve Cave (3), *E. latitans* from Honey Creek Cave (n=3), *E. latitans* from Honey Creek State Natural Area (3), *E. pterophila* from Comal Springs (3), *E. neotenes* (3), *E. troglodytes* (1), *E. waterlooensis* (n=1), *E. naufragia* (n=1), *E. tonkawae* from Testudo Tube (n=1), *E. tonkawae* from Bull Creek (n=1), *E. chisholmensis* (n=1), *E. sp*. from Georgetown Spring (n=1), *E. sp*. from New Braunfels well (n=3), and *E. sp*. 1, 2, and 3 as identified in (25) (n=1 each). We recognize the relatively low support for the genetic distinctiveness of three species (*E. neotenes*, *E. latitans*, and *E. pterophila*), and there is ongoing work to resolve this (105). However, the focus of this manuscript is not to determine taxonomic status; therefore, we adopt the current taxonomy (25). Salamanders were euthanized using 0.2% MS222 and decapitated in preparation for fixation and diceCT scanning.

We examined eyes from adults of six species (*Eurycea nana*, *E. sosorum*, *E. pterophila* (Comal Springs, CS), *E. pterophila* (Preserve Cave, PC), *E. latitans* (Honey Creek Cave, HC), and *E. rathbuni*) using diffusible iodine-based contrast-enhanced computed tomography (diceCT; (106-107, *SI Appendix*, Methods) and optimized at reconstruction (108, *SI Appendix*, Methods) to reveal soft tissues. We divided our sample into two ecotypes. Three species (*E. latitans, E. rathbuni,* and *E. pterophila* PC) exhibit subterranean morphology, and three (*E. nana, E. sosorum*, and *E. pterophila* CS) exhibit surface morphology. We took the relative eye volume (mm^2^) by measuring total volume of functional eye tissue, defined as the sum of lens and retina volumes, and standardized by gular fold to snout length (GFL). We performed a generalized linear model ANOVA and Tukey’s post (109) hoc for each developmental stage from 1–5 months post-oviposition (mpo) and adult for five species, one of which was represented by two ecotypes.

Details of fixation, staining, and CT Scanning, rendering, segmentation, and immunohistochemistry are presented in the more details in *SI Appendix*, Methods.

## Acknowledgments

We thank Katie Bockrath, David Britton, Desiree More, Braden West, Justin Crow, Adam Daw and other staff of the San Marcos Aquatic Resources Center, United States Fish and Wildlife Service for providing the salamander larvae that made much of this work possible. We also thank; Debbie and Don Davis, Ryan Hoffman, Zach Adcock, Pete Diaz, The Bexar Grotto Group, Staff at Guadalupe River State Park, Honey Creek State Natural Area, and Lost Maples State Natural Area, Dee Ann Chamberlain, Rustin Tabor, James Peterson, Jacob Owen, Marco Jones, Shannon Carrasco for assistance with site access and collecting. Core facility staff Jessie Maisano, Matthew Colbert, David Edey, Alissa Savage, Jacob Bisbal, and Jacob Armitage for training and guidance.

Guillaume Dury for help with navigating Inkscape and constructive critique of the figures through several iterations, and Christian Teague for help with westerns. The equipment in the ARSC is supported by funds from the Materials Applications Research Center of Texas State University. This work was supported by the National Science Foundation grant DEB2032632 (DMH and TJD) and DEB203263 (DMG); NSF had no involvement in the conduct of the study, preparation of the manuscript, or decision to submit it for publication.

