## Supplemental Appendix for "Parallel Sensory Compensation following Independent Subterranean Colonization by Groundwater Salamanders (*Eurycea*)"

### Supplementary Material

#### Methods

##### Phylogenetic and Comparative Analyses

For discrete traits, we estimated four Mk models (43) of character state evolution using phytools::fitMk() and three phenotypes (Biphasic, Surface, and Subterranean): one-rate, six-rate (all rates different), ordered (Biphasic ↔ Surface ↔ Subterranean, four rates) and irreversible (Biphasic → Surface → Subterranean, two rates). We repeated this using phytools::fitgamma Mk() to account for rate heterogeneity in state changes across the branches.

To quantity the rate of evolutionary shifts among the three ecotypes (Biphasic, Surface, and Subterranean), we used ancestral reconstruction using four Mk models (43): one rate (all rates equal), six rate (all rates unconstrained), ordered (Biphasic ↔ Surface ↔ Subterranean, four rates), and irreversible (Biphasic → Surface → Subterranean, two rates). Because we suspected that rate change across the tree might be heterogeneous, we repeated these four analyses using phytools::fitgammaMk(), which estimates an additional gamma parameter describing the distribution of rates along the edges. For all analyses, we fixed the state of the root as biphasic because a biphasic lifecycle is ancestral for this clade (42).

##### Stochastic Character Mapping

Stochastic character mapping (45), a Markov Chain Monte Carlo procedure that simulates a phylogenetic map of history of a discrete character under a specified model, was done using phytools::make.simmap(). Following a burn in of 2000 generations, 10,000 total generations were sampled at a frequency of 10, yielding 1000 posterior samples of character maps; these were summarized to yield posterior probabilities of ancestral states and statistics describing the frequency of character state transitions. We analyzed the ecotype data under four Mk models and four corresponding Mk gamma models; the latter include a rate heterogeneity parameter. We excluded *E. wallacei* from the analysis because other *Eurycea* species in this region of the phylogeny are poorly sampled, and inclusion of *E. wallacei* which has extreme values, seemed to bias the trait estimates of the deepest nodes.

We compared the eight models using the Akaike Information Criterion (*SI Appendix*, Table S1). The best model (one-rate gamma) had an AIC weight (AICw) of 0.435, followed by one-rate, ordered, and ordered gamma (AIC weights = 0.204, 0.151, and 0.131), in total accounting for 92% of the weight. However, AIC values of these models do not differ much, and likelihood-ratio tests between most model pairs do not reject the null hypothesis. Given that no one model overwhelmingly dominated, we used a weighted average of all models (44), rather than a single model, to estimate ancestral states. Because it is not possible to calculate ancestral states under the weighted model, we also estimated ancestral states (*SI Appendix*, Fig. S3) under the one-rate-gamma model, the best model. The marginal (empirical Bayesian) probabilities of each state are shown as pie charts at the internal nodes in Fig. 1. The ancestral reconstructions of most nodes under this model were not decisive; for many internal nodes, each ecotype had roughly the same probability.

Under likelihood, the uncertainty in ancestral state estimates precludes a direct count of the numbers and types of state changes. To do this, we obtained estimates of the numbers of state changes and confidence intervals using stochastic character mapping (45) by simulating state changes under the model and summarizing these over all simulations.

As an example, we chose the ordered $\Gamma$ model. Under this model (*SI Appendix*, Table S2), there were no changes from biphasic to subterranean or the reverse (a constraint of the ordered model, which requires an intermediate step through the surface state). The median number of changes from biphasic to surface was three (95% HPD, 2–7), two (1–6) from surface to biphasic, seven (6–12) from surface to subterranean, and three (0–10) from subterranean to surface. The large 95% HPD intervals for some categories reflects the uncertainty of the ancestral state reconstructions, as well as the extensive homoplasy in transitions between surface and subterranean ecotypes. Estimates under other models are included in *SI Appendix*, Table S2, and all indicate large numbers of reversals and convergences.

#### Fixation, Staining, and CT Scanning

Specimens were collected, euthanized, and immediately fixed overnight using PAXgene Tissue FIX (Qiagen, PreAnalytics, Cat. No. 765312), washed fifteen minutes in PAXgene Stabilizing Solution (Qiagen, PreAnalytics, Cat. No. 765512), and placed in fresh stabilizing solution before storage at -80º C. Specimens remained in -80º C until staining. As a variation on the standard diceCT protocol, Green et al. (106) demonstrated that an iodine-contrast agent using PAXgene Stabilizing Solution as the base solvent (e.g., instead of ethanol) preserves both biomolecules and morphologies. We mimicked this approach by preparing a 2% weight-to-volume mixture of PAXgene Stabilizing Solution and iodine metal (106, 107) to stain most of our samples (i.e., those older than one-month post-oviposition). For specimens just one-month post oviposition, which are especially small, we instead prepared a lower-concentration, 1% weight-to-volume staining solution. As with high concentrations of ethanol (106), iodine crystals dissolve readily into Paxgene Stabilizing Solution. To ensure complete dissolution, the solute was thoroughly mixed by hand and left overnight before first use (106).

All tissues were imaged at The University of Texas High-Resolution X-ray Computed Tomography Facility (UTCT) and were scanned at room temperature using an Xradia MicroXCT 400 (Zeiss). Because biological specimens are expected to have slight variation in staining, each specimen was individually optimized at reconstruction to utilize the full 16-bit dynamic range of the detector (108). Scan settings for heads representing the developmental series were as follows: 4X objective, 70 kilovolts (kV), 8.5 watts (W), 0.25–0.5 second (s) acquisition time, detector position at 10.002 millimeter (mm). Scan settings for heads representing the adult stage were as follows: flat panel, 70 kV, 8.5 W, 0.06 s, 5 frames per view, detector position at 155.707 mm. Voxels were isometric, ranging from 4.00 x 4.00 x 4.00 micrometers (µm) for specimens in the developmental series to 7.00 x 7.00 x 7.00 µm for adults. Following scanning specimens were returned to a 15 mL collection tube filled with fresh PAXgene Stabilizing Solution and stored in -80º C where the iodine was allowed to passively diffuse.

#### Rendering and Segmentation

DiceCT data was reconstructed by Xradia Reconstructor. Sixteen-bit Tagged Image File Format (TIFF) series were imported, rendered, and soft tissues segmented using Dragonfly software, version 2020.2 (Object Research Systems, ORS Inc.). Soft-tissue segmentation of the lens and retina was accomplished using the magic wand feature, then detailed using the automated tool, finally the volumes for each segmentation were recorded. We mapped and quantified neuromasts of the ALL of salamanders from seven species of *Eurycea* (four surface and three subterranean species; Fig. 4), using diceCT scans of three individuals per species. Neuromasts were identified using the sphere-segmentation feature placed over each neuromast. Each respective region (see Fig. 4) was colored differently and hand counted for downstream analysis. Both datasets were analyzed using R Statistical Software (109).

#### Immunohistochemistry and Imaging

Specimens were euthanized and fixed overnight, using 4% paraformaldehyde prepared in PBS. Then they were washed three times in PBS for 15 minutes/wash and incubated overnight at 4ºC in 30% sucrose to cryoprotect the tissue during sectioning. The sucrose solution was prepared in PBS with 0.1% sodium azide as an antimicrobial agent. We considered specimens to be cryoprotected once they sank to the bottom of the tube. Specimens were embedded in TissueTek® and cut into 15 µm sections at -22º C using a Leica Cryostat equipped with a disposable blade. Sections were collected on gelatin-coated glass slides and stored at -20º C until use.

Immunolabeling was accomplished as follows: Sections were incubated and blocked for one hour at room temperature in PBS containing 0.3% Triton X-100 (LabChem, LC262801) and 5% non-fat dry milk as a blocking agent. Following blocking, sections were washed three times for 10 minutes/wash with PBS. Sections were then incubated with anti-PAX6 antibody (Thermo Fisher/Invitrogen, PA1-801) diluted 1:75 in PBST containing 0.1% non-fat dry milk. Negative controls were not incubated with anti-PAX6, but with PBST containing 0.1% non-fat dry milk. Following a two-hour incubation at room temperature, slides were washed three times for 10 minutes/wash in PBS. Sections were then incubated for one hour at room temperature with goat anti-rabbit secondary antibody conjugated to Cy5 (Thermo Fisher Scientific/Invitrogen, A10523), diluted 1:500 in PBST containing 0.1% non-fat dry milk. Sections were then washed three times for 10 minutes/wash in PBS. Coverslips were mounted onto the slides with EverbriteTMFluorescence Antifade Mounting Medium (Quartzy/Biotium, 23002) containing DAPI as a nuclear stain. Slides were sealed with fingernail polish and stored frozen at -20º C until imaged.

Images were obtained using an Olympus FV1000 Laser Scanning Confocal Microscope. Three neuromasts from each of three individuals of each of the two species were imaged. Confocal laser settings were optimized for the DAPI channel for each section. The laser settings for the Cy5 channel were optimized using a sample generated from a 1mpo Texas blind salamander, and the same Cy5-settings were used for all images acquired. Images were captured as Z-stacks comprising 30 optical sections with a step size of 0.41 µm and were prepared for publication using Adobe Photoshop 24.5.0. The fluorescence intensity of PAX6 labeling within the boundaries of the mantle cells was quantified using Olympus's Fluoview Fluorescence Intensity Analysis Tool. The maximum fluorescence intensity from each Z plane was summed for each neuromast, and that sum was averaged among the three neuromasts for each individual. For the negative controls, the sum of the maximum fluorescence intensity for one neuromast was used to reference but not included in statistical analysis.

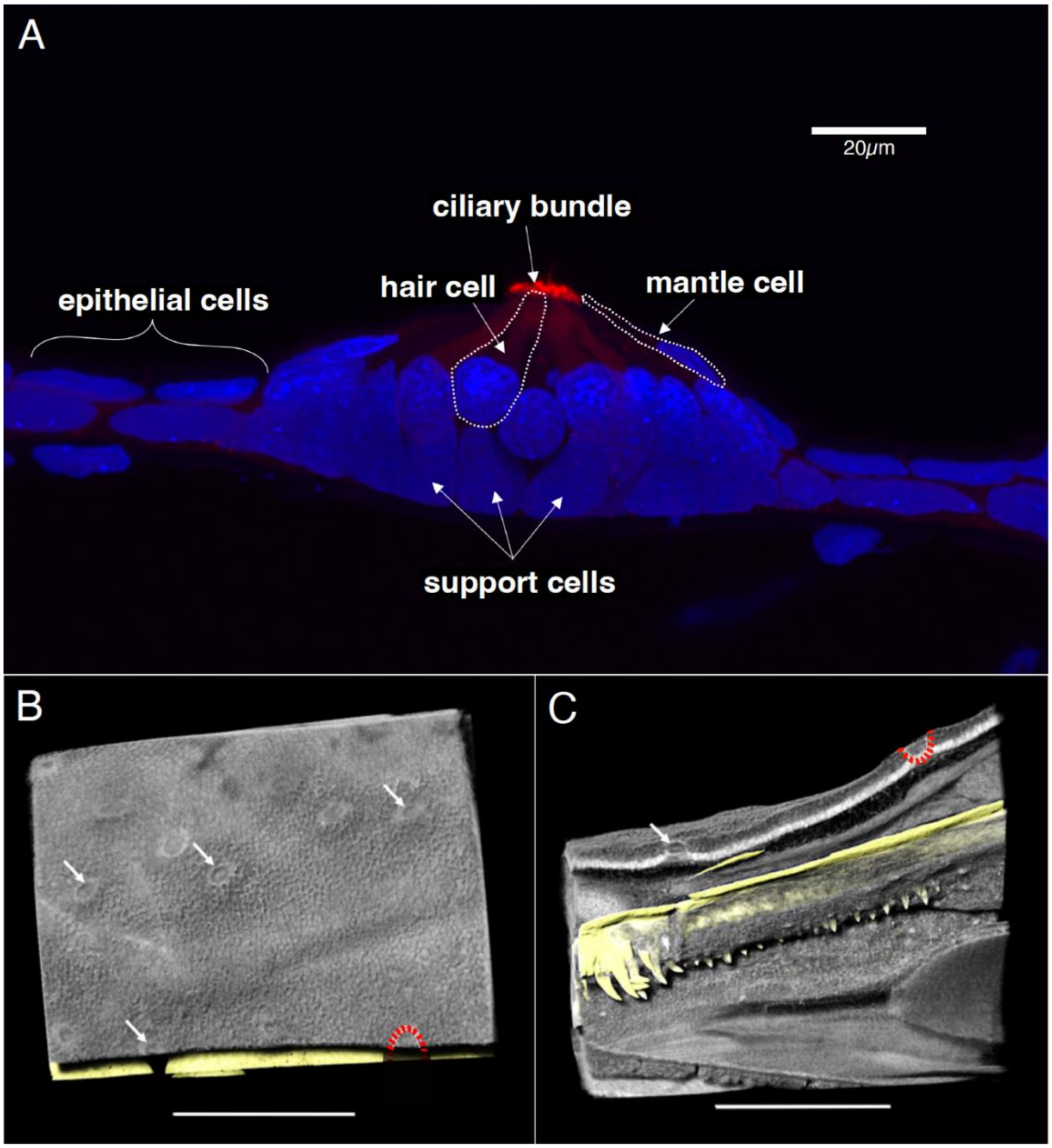


**Supp Figure 1.** Neuromast identification in confocal image (A) and diceCT scans (B and C). Cellular components of the neuromast are stained with DAPI (blue) and labeled with anti-PAX6 (red) and can be observed in panel A; the mantle, support, and hair cells are identified by morphology and position within the epithelial layer (Scale bar is 20 µm). Neuromasts are observed on the surface of an adult *E. rathbuni* (B–C). A diceCT scan at greater magnification shows several neuromasts with typical round morphology on the surface of the epidermis (B, white arrows and red dashed lines mark outlining). A cross-section (C) shows the z-plane of the neuromast (white arrows and dashed line mark outlining). Scale bars in B and C are 1 mm.

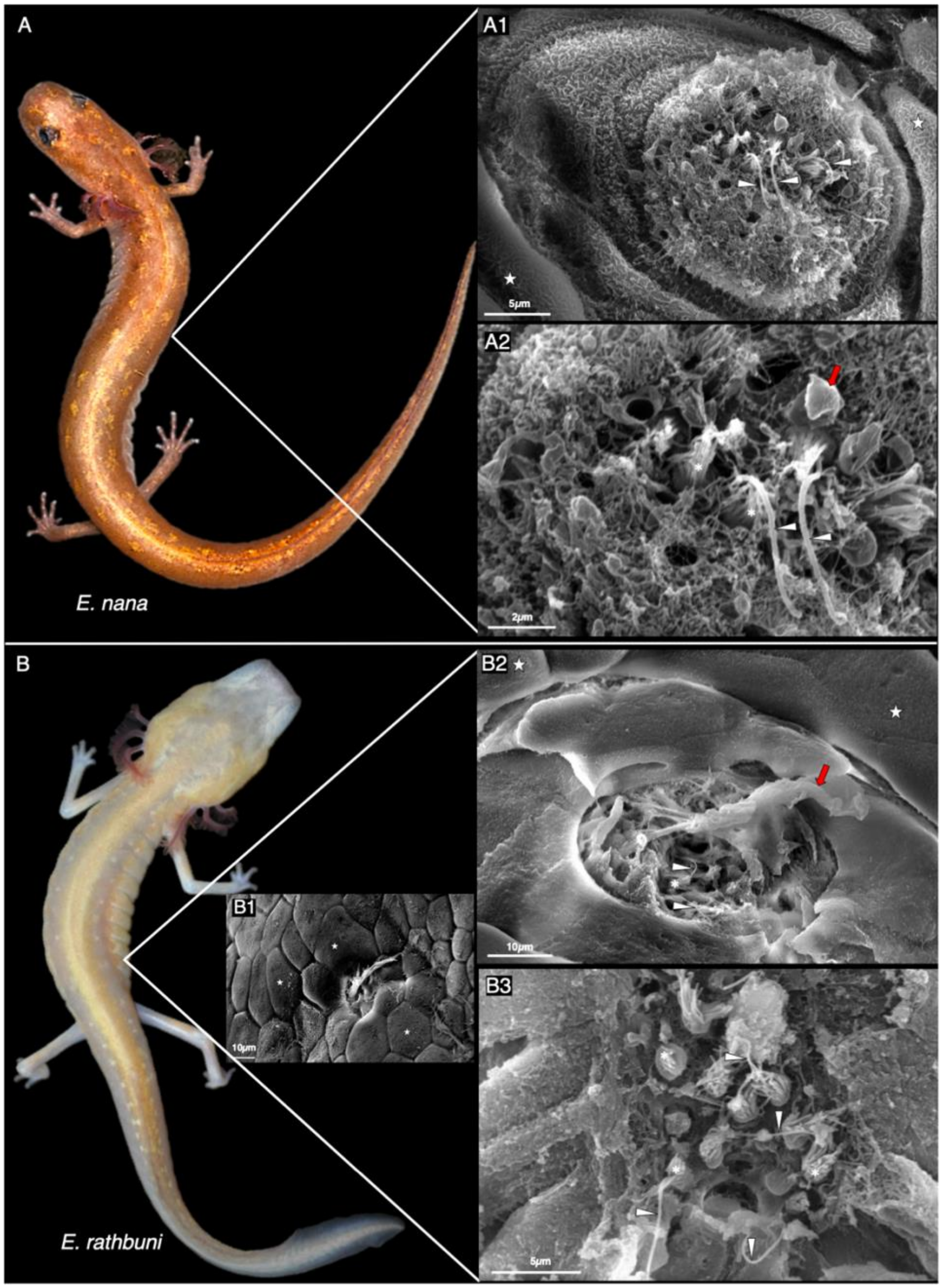


**Supp Figure 2.** The SEM of neuromasts in the surface species *E. nana* (A: 1-2) and the subterranean species *E. rathbuni* (B1 B2). Two specimen images are represented for each; the first for each species shows an entire neuromast with epidermal layer (stars) and cupula (red arrow) (A1 and B1-2), and the second shows higher magnification of neuroast ultrastructure, including kinocilia (traingles) and stereocilia (asterisks) (A:2 and B:3). We identify a pit organ found in *E. rathbuni* (B1). Scales bars are shown in each micrograph.

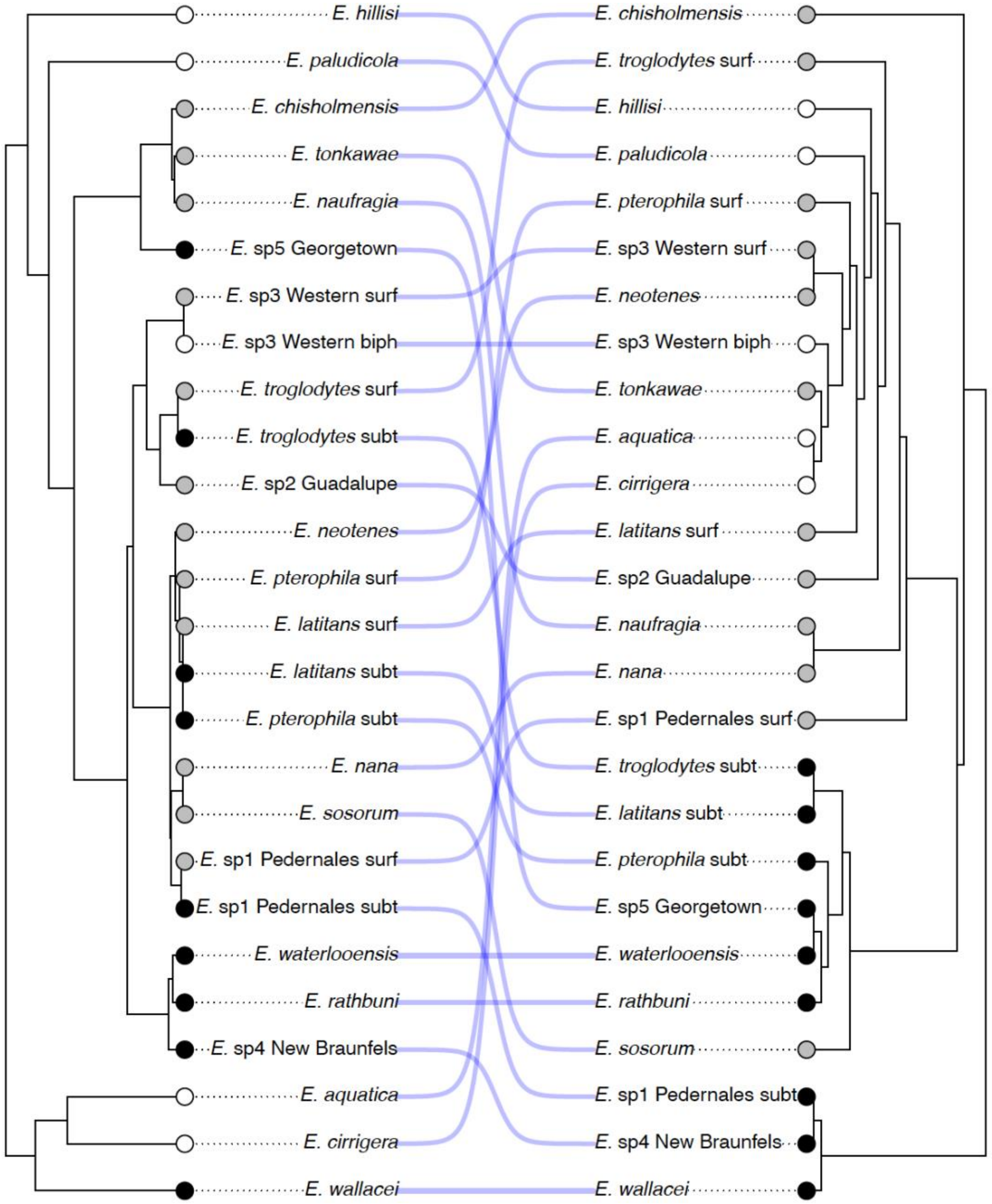


**Supp Figure 3.** "Tanglegram" showing tip states of life history mapped onto the DNA phylogeny (left) and a minimum evolution tree (right) estimated from three variables (Retina Total Volume, Lens Total Volume, and SGL) using euclidean distance. The tree branches are rotated to minimize the number of line crossings. The normalized Robinson-Foulds distance between the two trees is 0.958, indicating that the topologies are highly dissimilar. Circle colors indicate (white = Biphasic, gray = Surface, black = Subterranean).

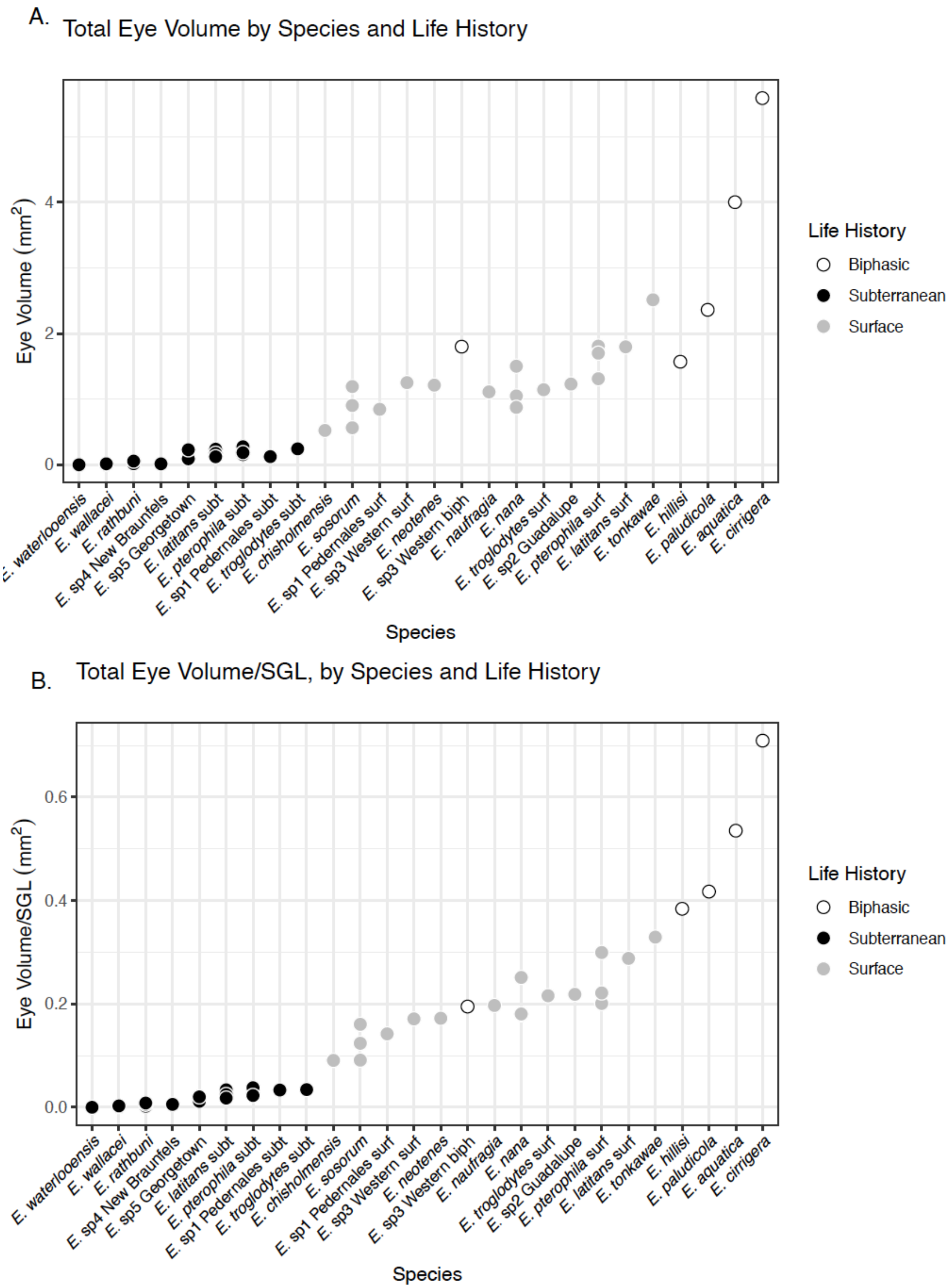


**Supp Figure 4.** Eye volume by species plots including, Eye Total Volume by species and life history (A) Eye Total Volume/snout to gular length (SGL), by species and life history (B). Each dot represents one individual.

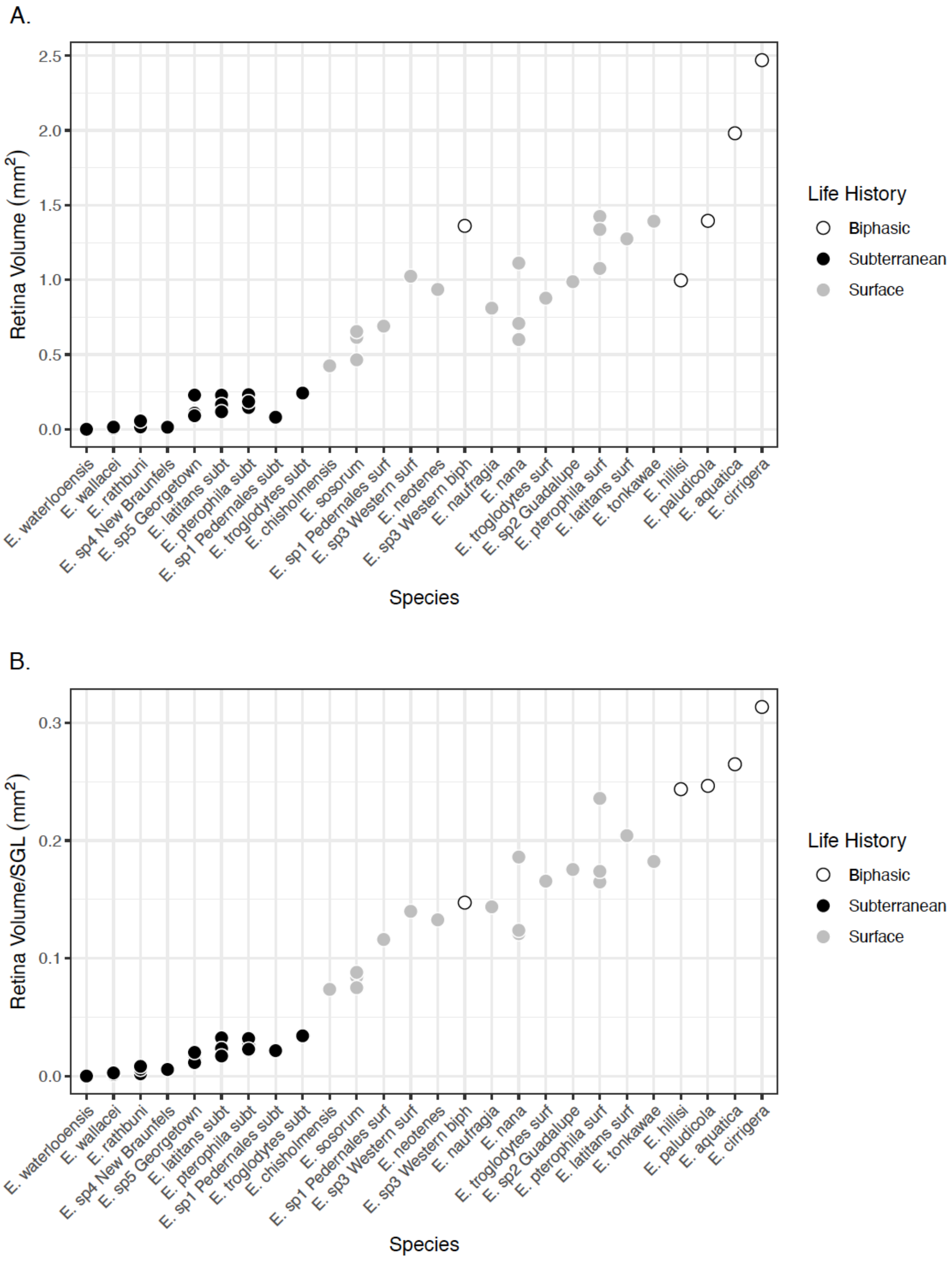


**Supp Figure 5.** Retina volume by species plots including, Retina Total Volume by species and life history (A) Retina Total Volume/snout to gular length (SGL), by species and life history (B). Each dot represents one individual.

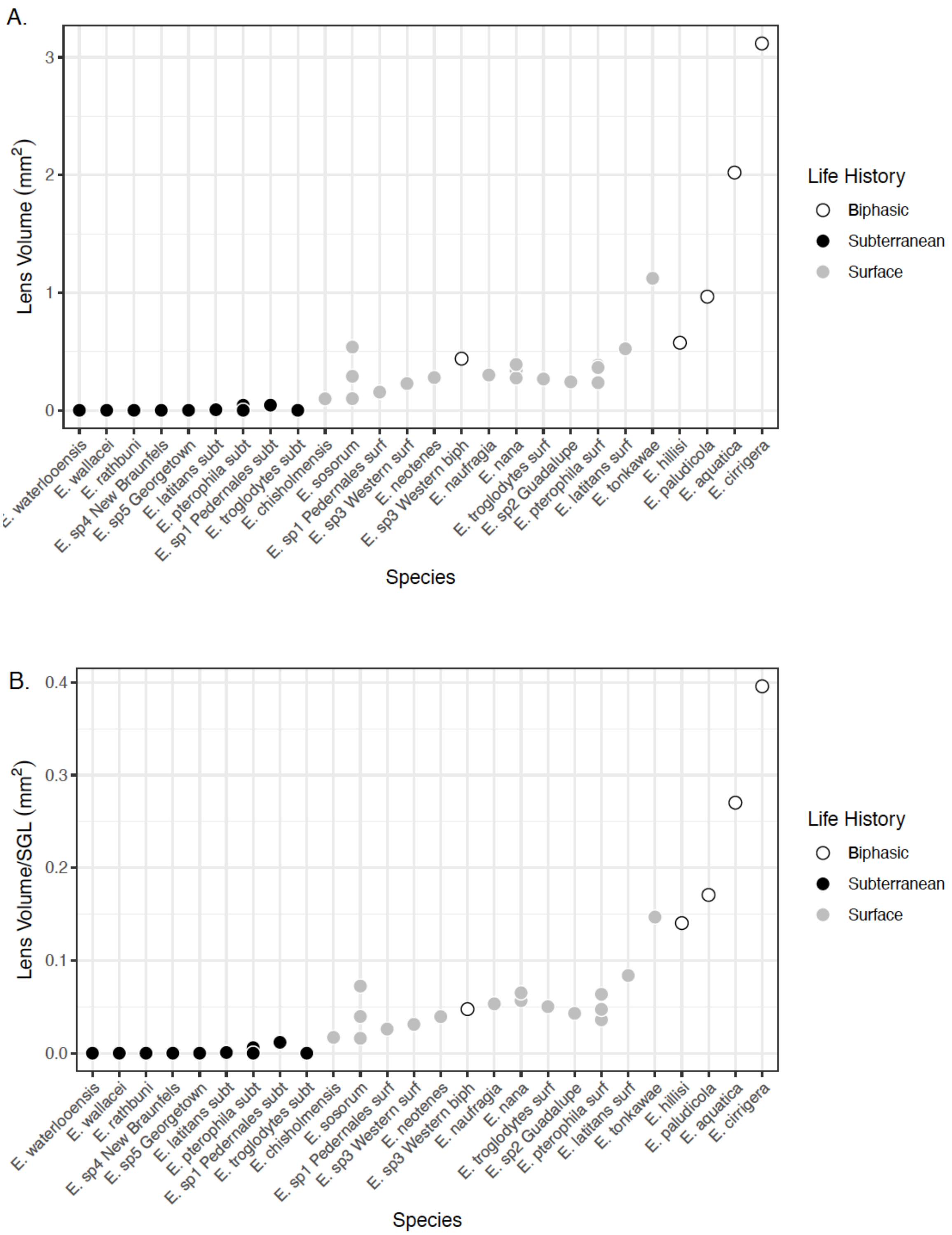


**Supp Figure 6.** Lens volume by species plots including, Lens Total volume by species and life history (A) Lens Total volume/snout to gular length (SGL), by species and life history (B). Each dot represents one individual.

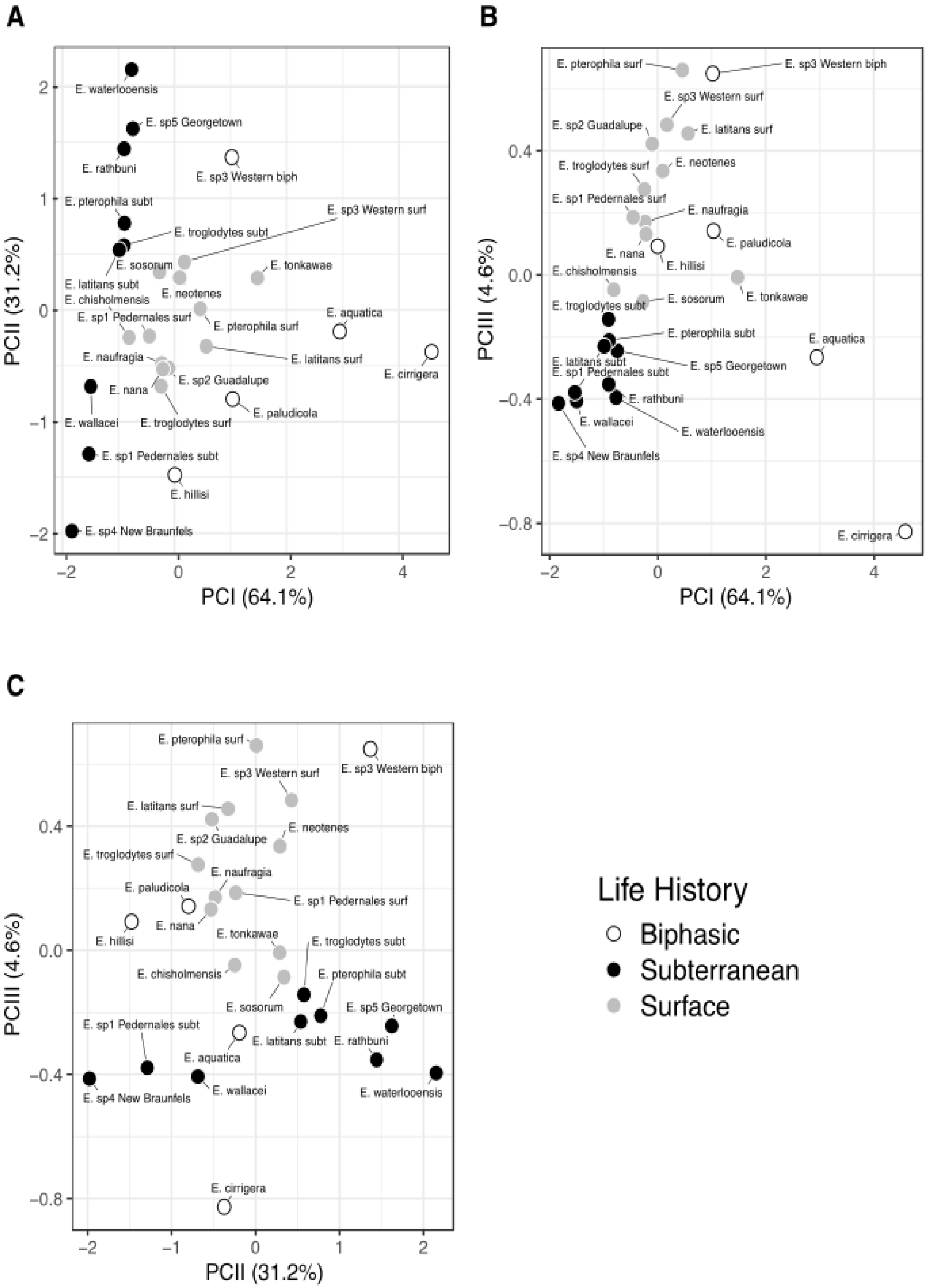


**Supp Figure 7.** Standard principal component plots calculated from Retina Total Volume, Lens Total Volume, and Snout-Gular Length using stats::prcomp(). Each point represents the mean of individual specimens within a species. The percent variation explained by each component is indicated on the graph axis.

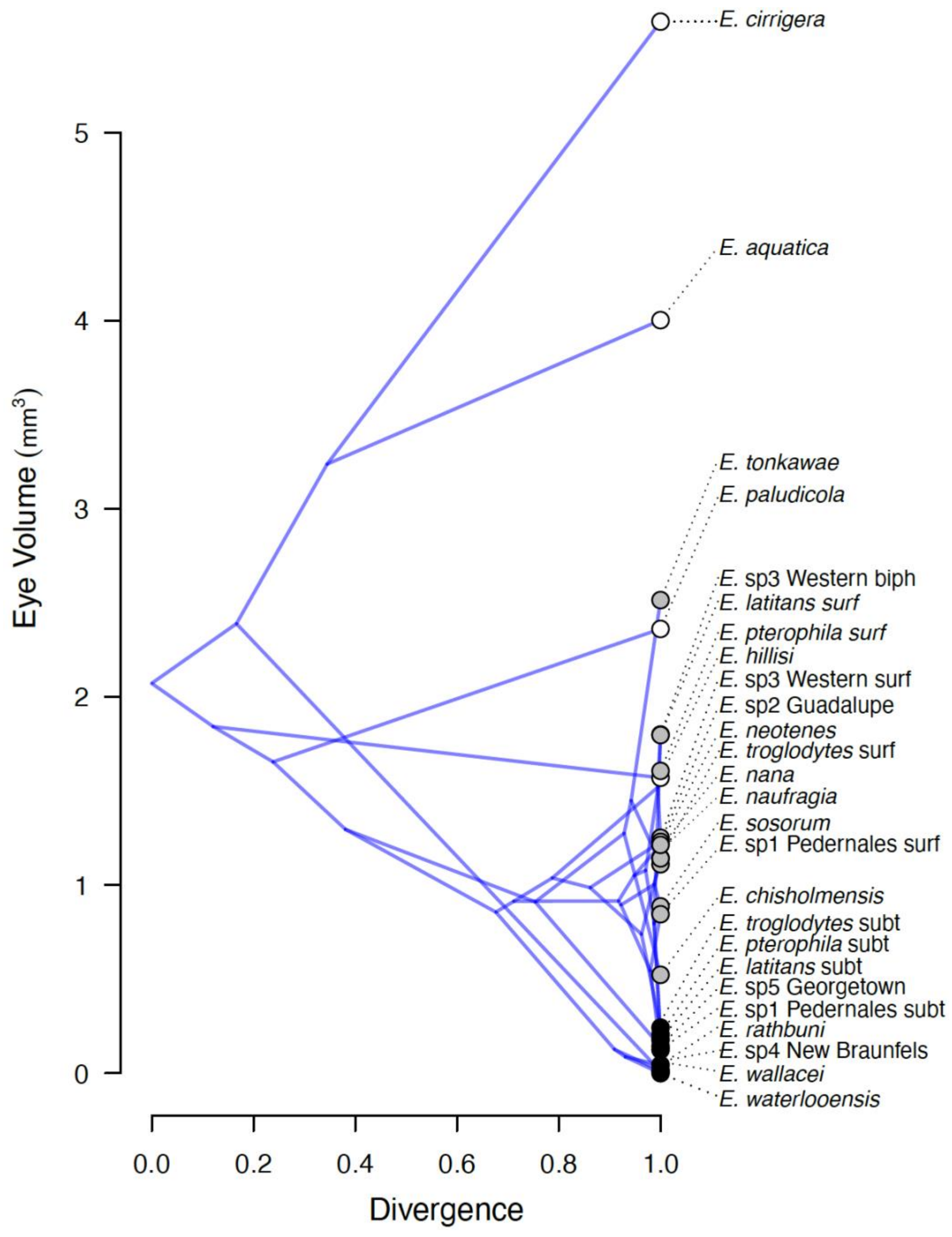


**Supp Figure 8.** Raw values for Eye volume (mm$^3$) by species (y-axis) plotted against scaled genetic divergence (x-axis) and superimposed on the chronogram, plotted by phytools::phenogram. Symbols are filled by life history (white = Biphasic, gray = Surface, black = Subterranean).

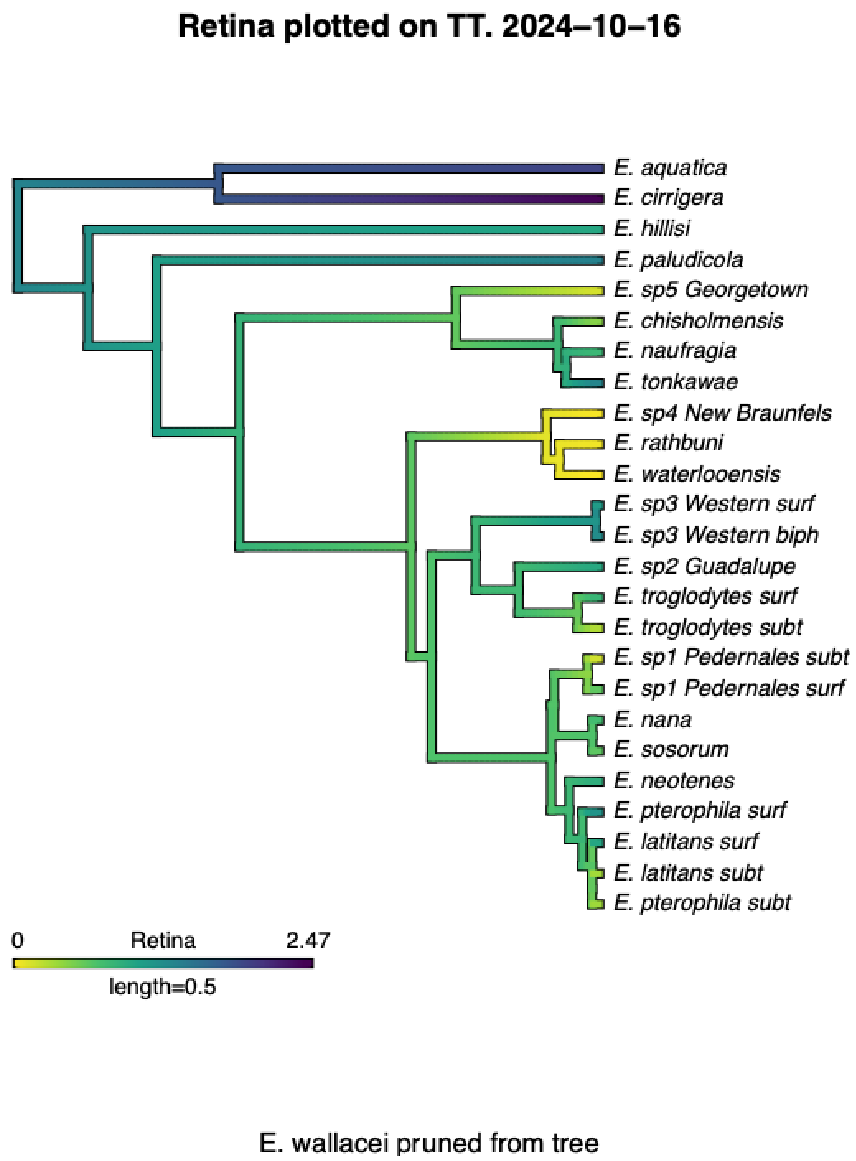


**Supp Figure 9.** Retina Volume plotted onto the phylogeny under a Brownian motion model.

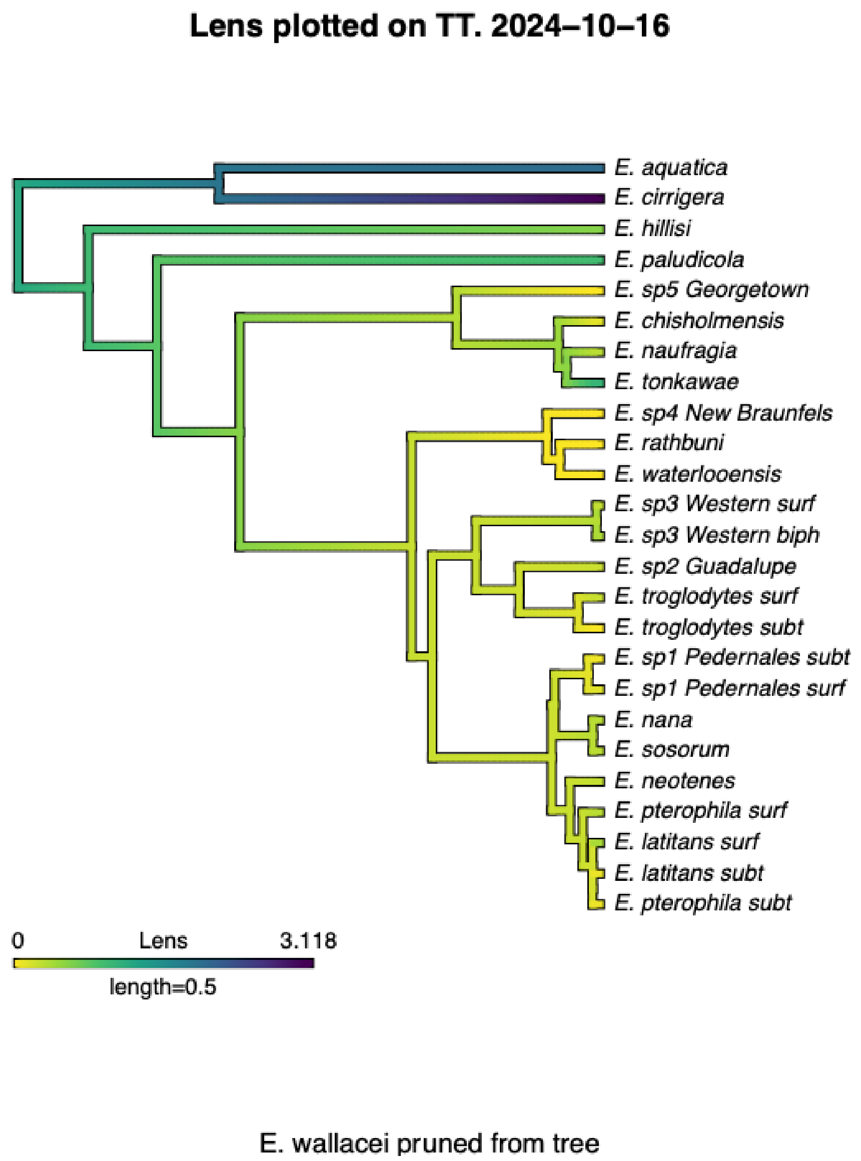


**Supp Figure 10.** Lens Volume plotted onto the phylogeny under a Brownian motion model.

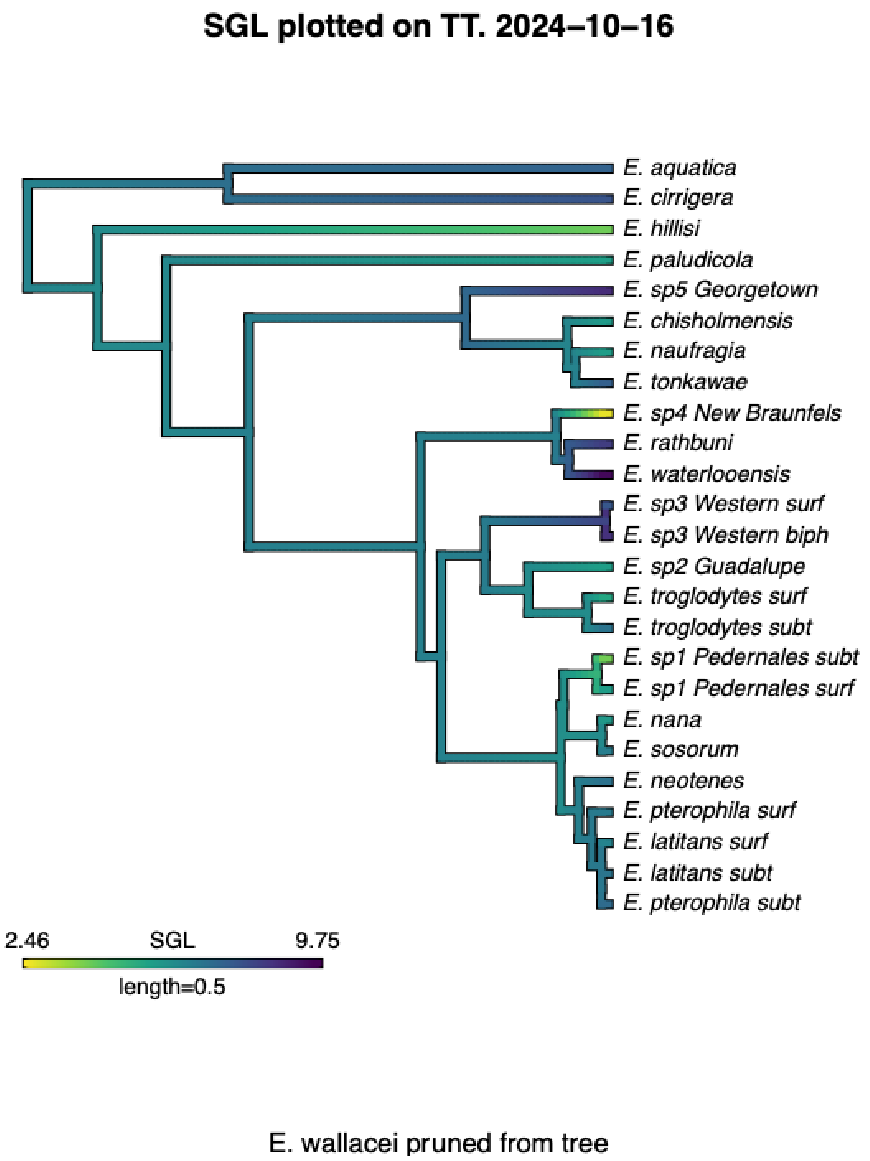


**Supp Figure 11.** SGL plotted onto the phylogeny under a Brownian motion model.

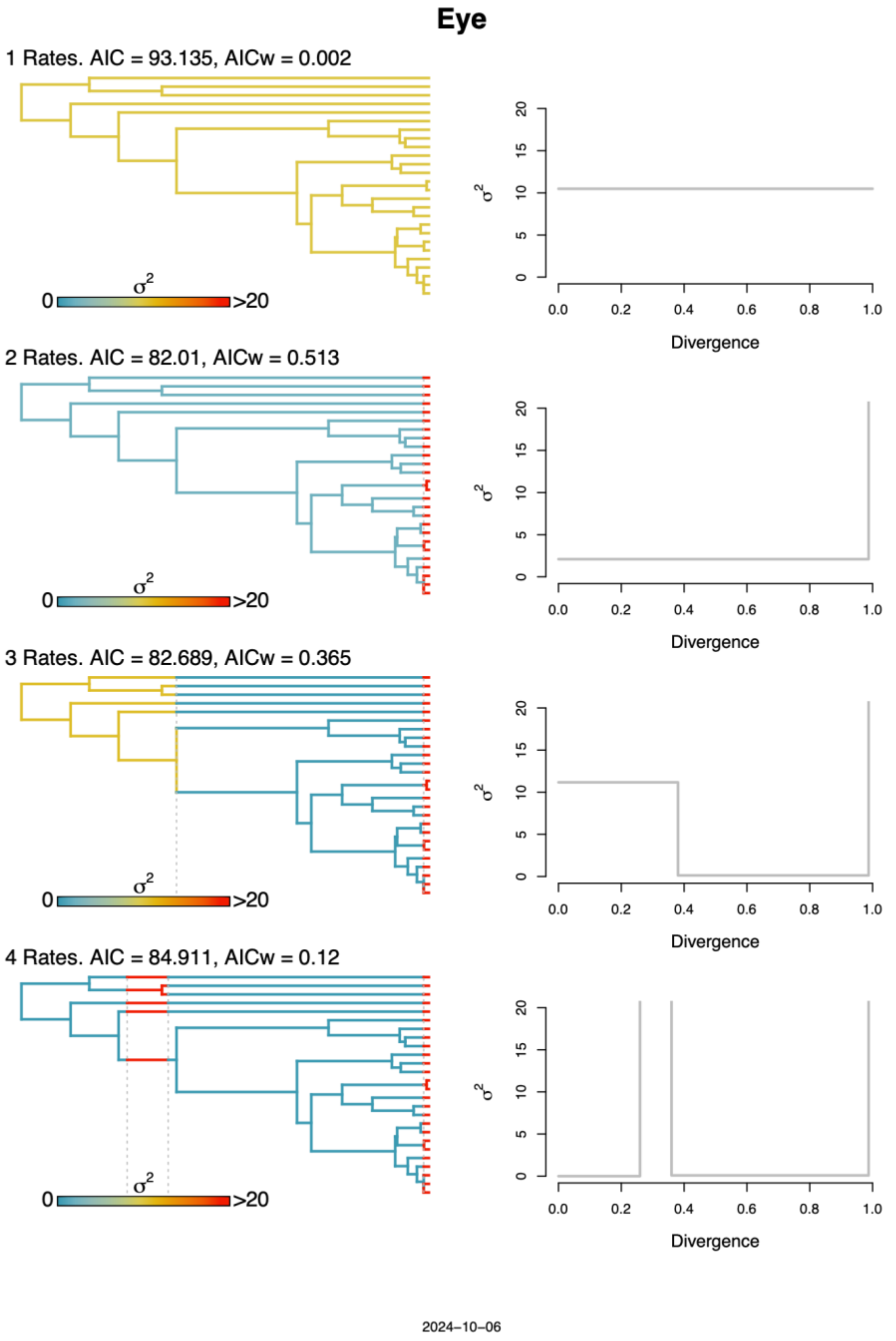


**Supp Figure 12.** Rate shift plots for Eye Volume. The best model was determined by the largest AIC weight.

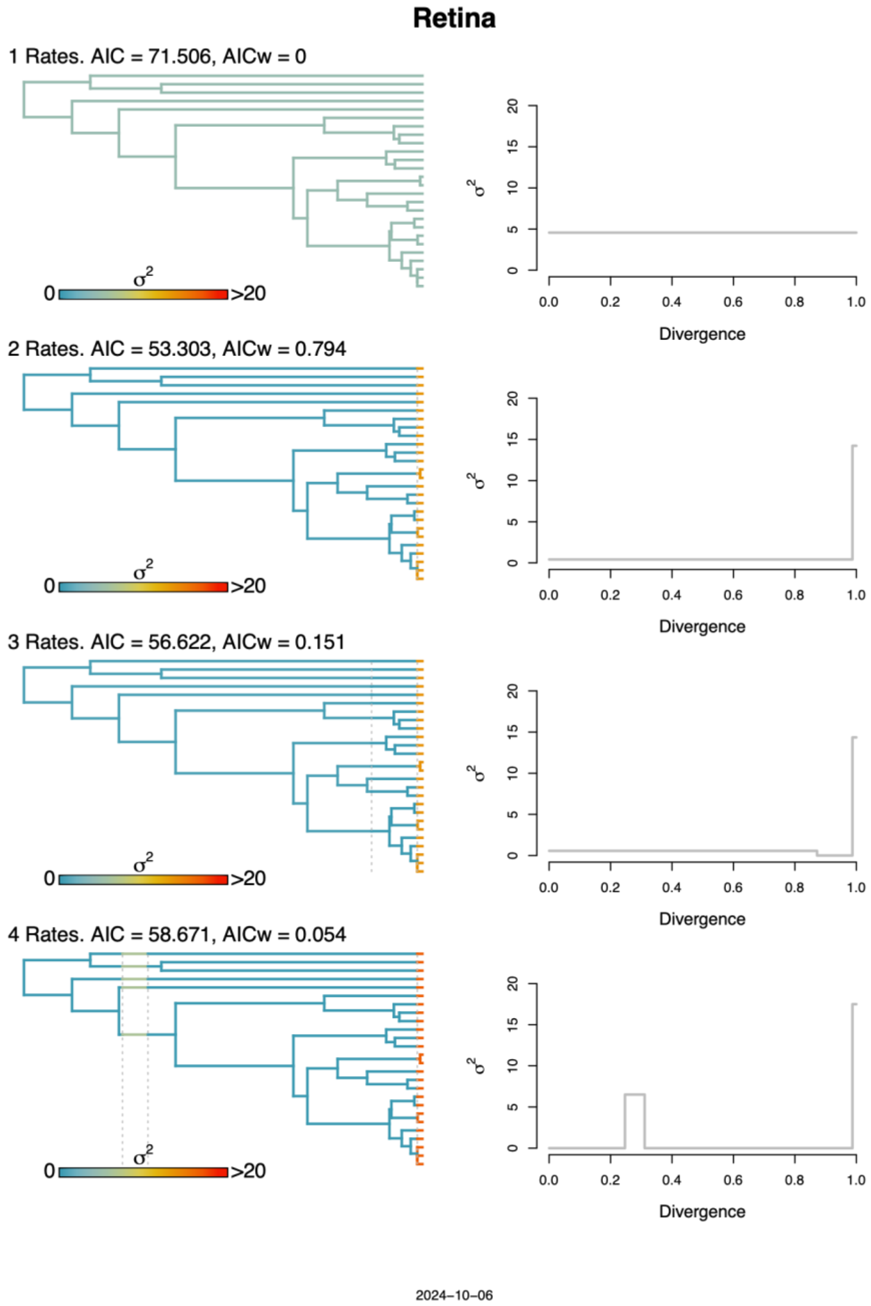


**Supp Figure 13.** Rate shift plots for Retina Volume. The best model was determined by the largest AIC weight.

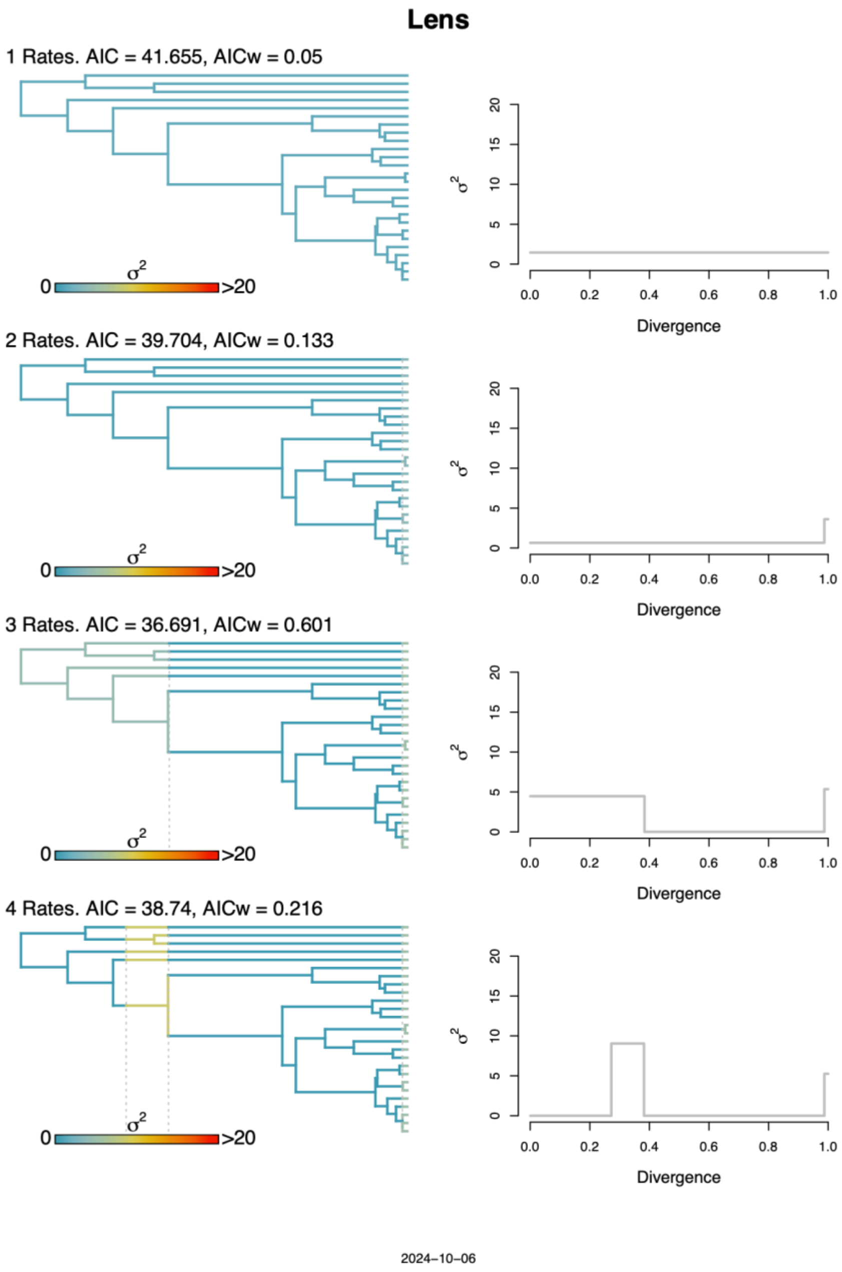


**Supp Figure 14.** Rate shift plots for Lens Volume. The best model was determined by the largest AIC weight.

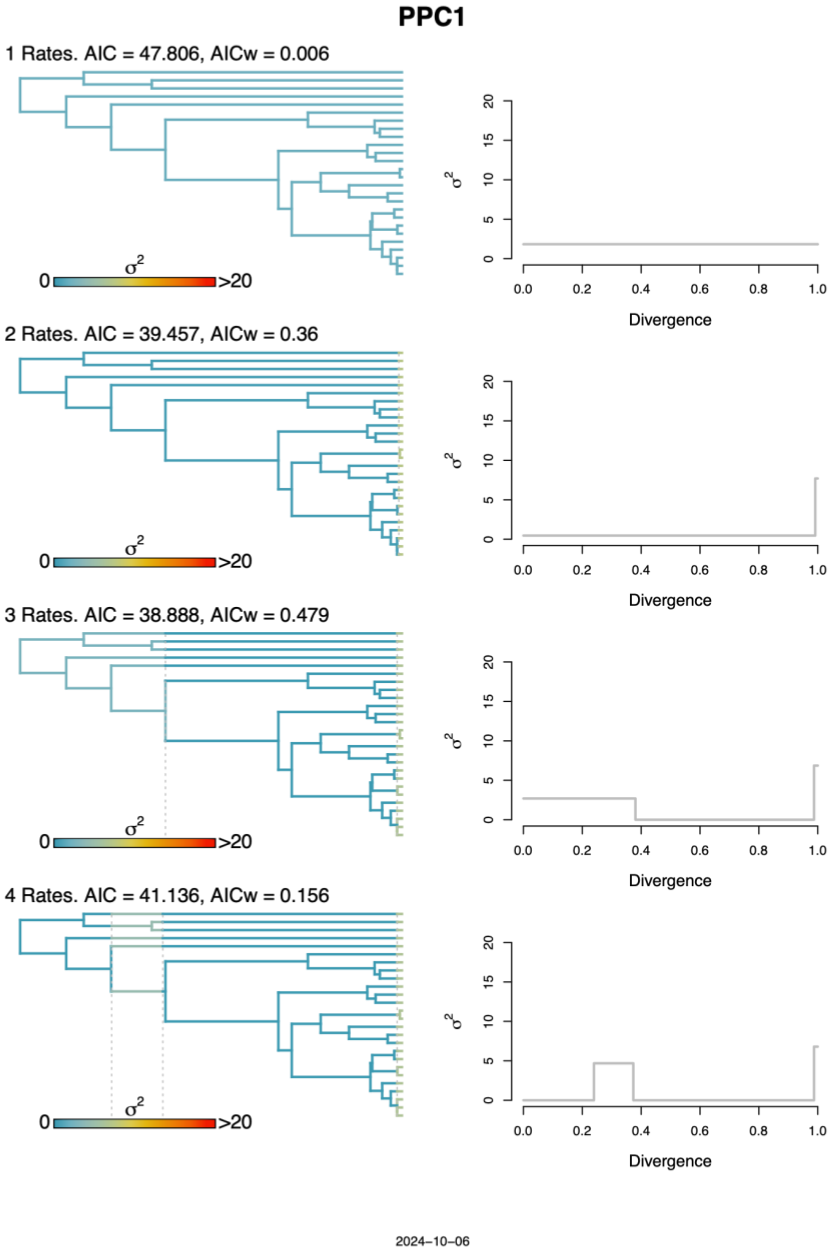


**Supp Figure 15.** Rate shift plots for PPCI scores. The best model was determined by the largest AIC weight.

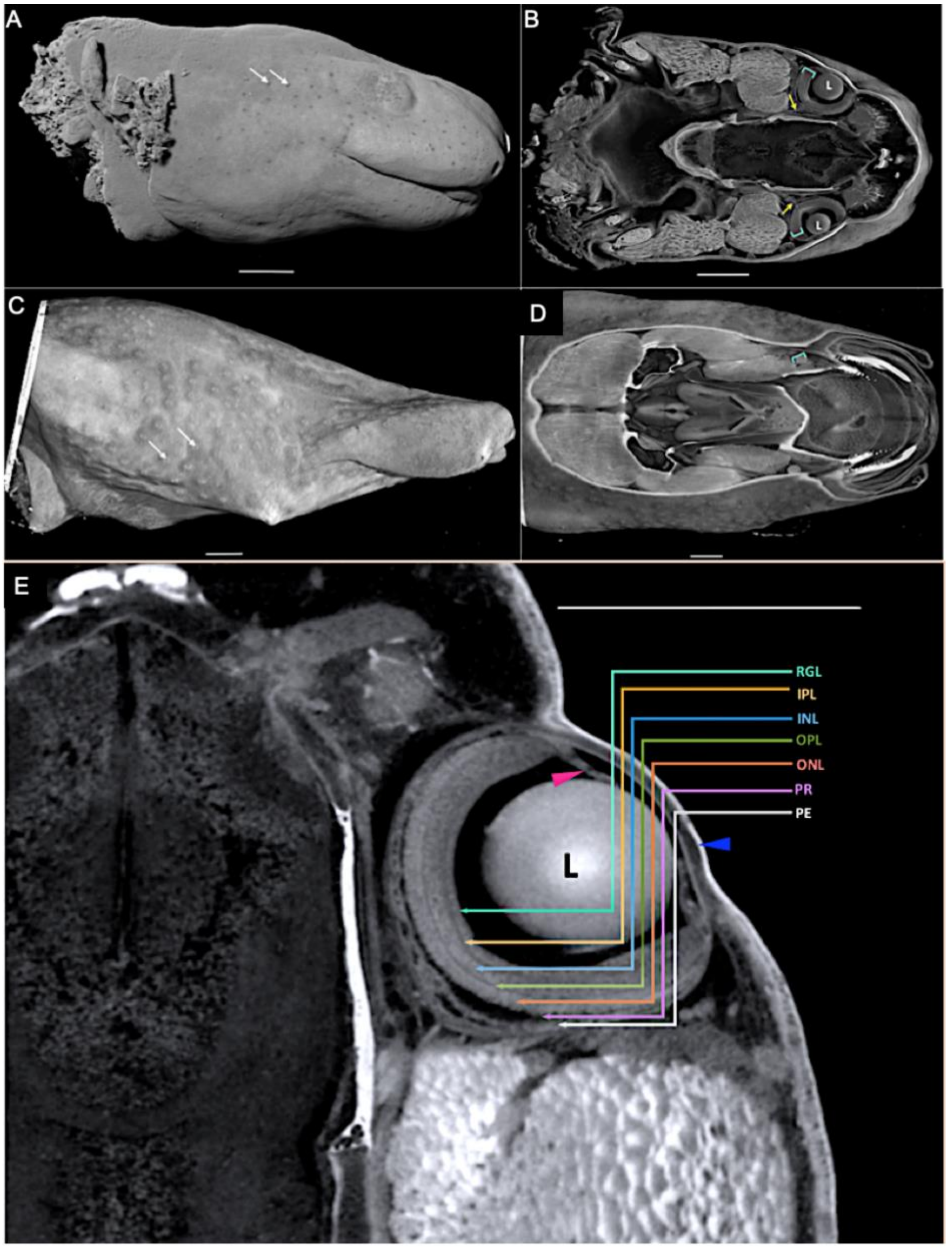


**Supp Figure 16.** Adult ocular structure observed using diceCT. In both *E. nana* (A) and *E. rathbuni* (C) surface neuromast structures can be observed (panel A and C, white arrows). Gross ocular anatomy is noted in *E. nana* 3D cross-section (B) including the lens (L), optic nerve (yellow arrows), and retina (green brackets). 3D cross-section of *E. rathbuni* (D) appears to have only an underdeveloped retina (green bracket). Several layers of the retina are noted in *E. nana* (E) including the retinal ganglion layer (RGL, green), inner plexiform layer (IPL, orange), inner nuclear layer (INL, blue), outer plexiform layer (OPL, green), outer nuclear layer (ONL, salmon), photoreceptor layer (PR, purple), and pigment epithelium (PE, light grey). In the same panel the lens (L), ciliary body (pink arrow), and cornea (blue arrow) are all noted. All scale bars indicate 1 mm.

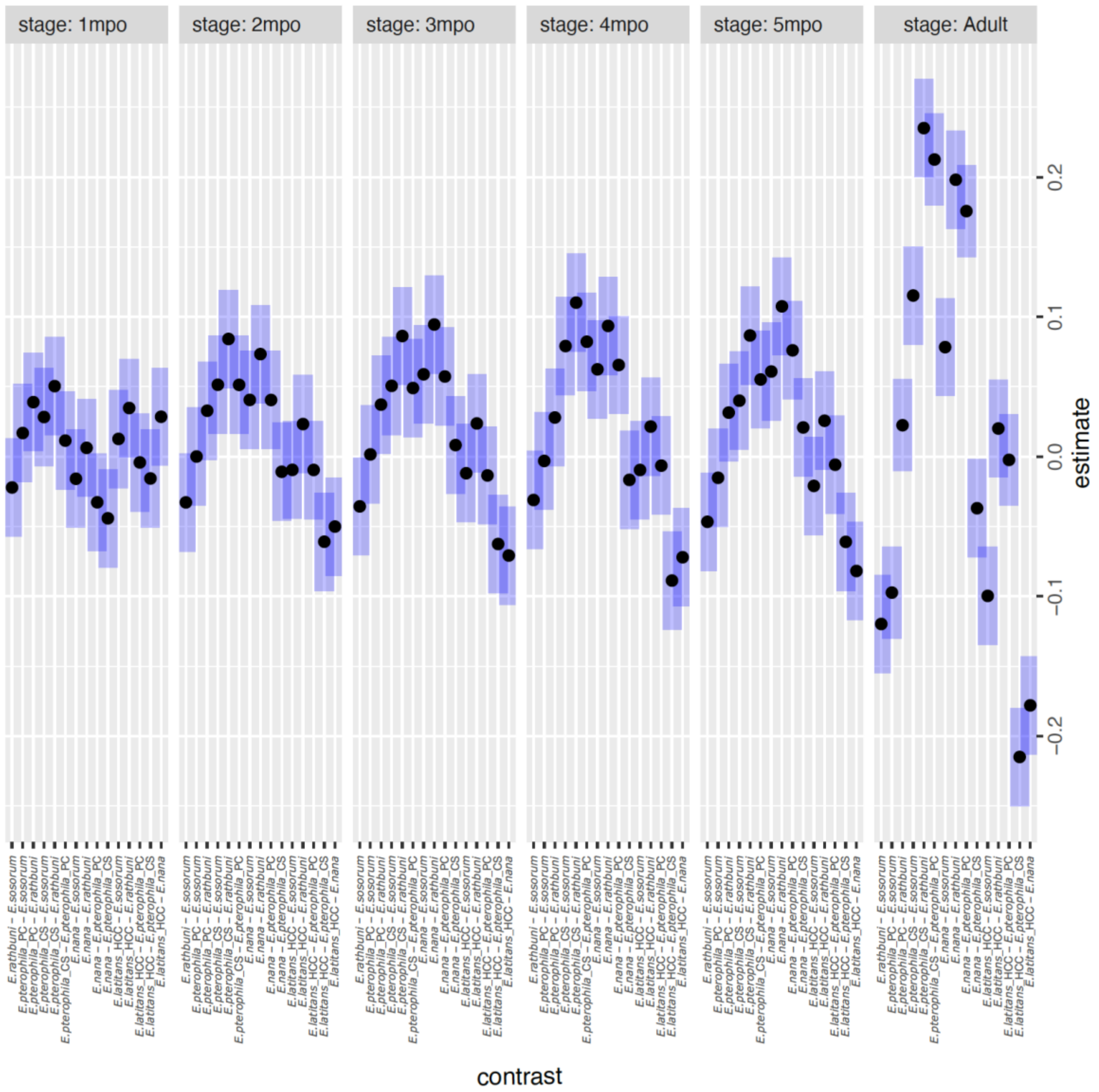


**Supp Figure 17.** Estimated marginal means for total eye volume through development. Six species are represented and marginal means are facetted by developmental stage 1-5mpo, and Adult.

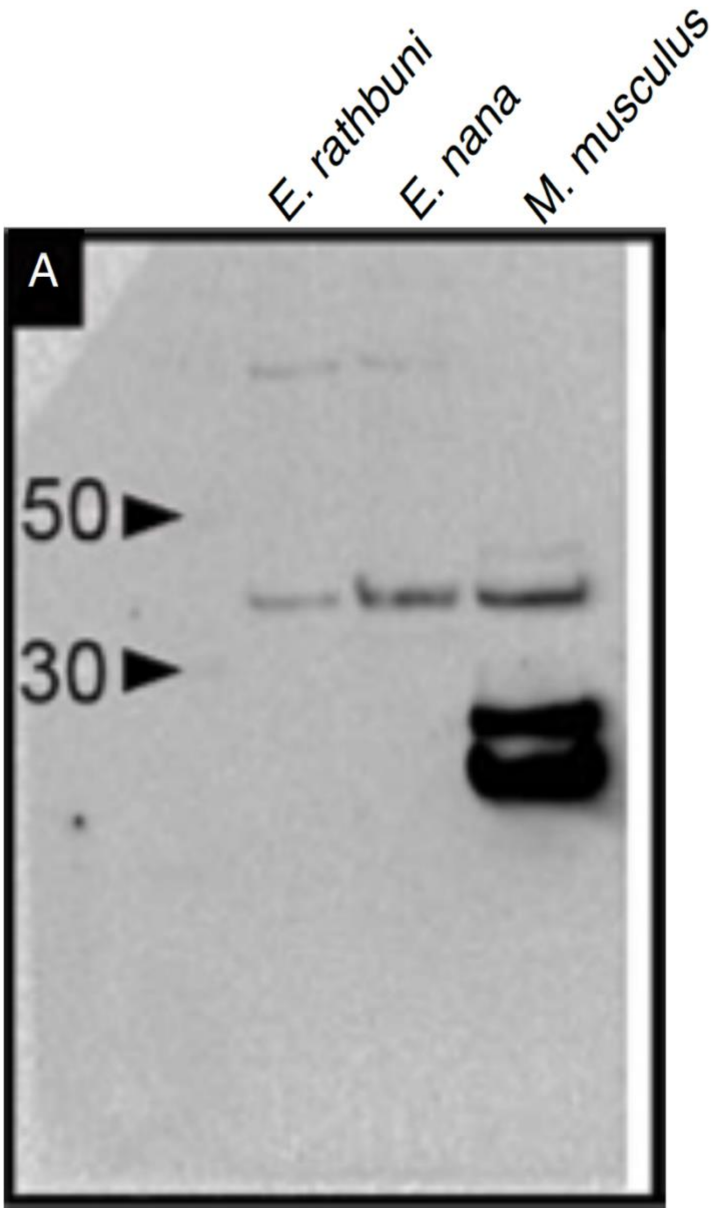


**Supp Figure 18.** PAX6 western blot. Lysates from embryos of two species of salamanders (*E. nana* and *E. rathbuni*) were used to perform western blot analysis of PAX6. A mouse (*M. musculus*) eye was included as a positive control. For all taxa, the labeled bands co-migrate.

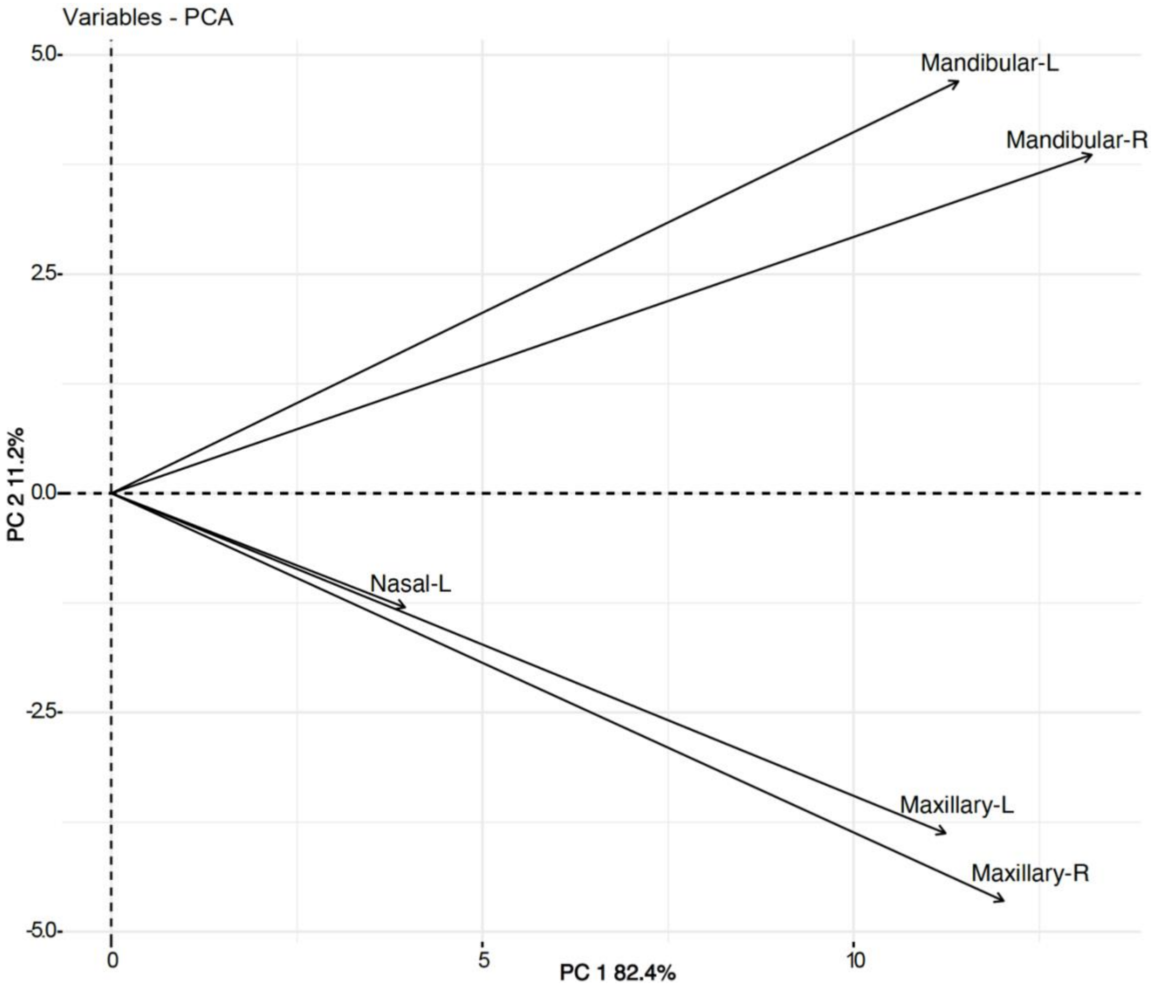


**Supp Figure 19.** Principal components analysis loadings for the respective neuromast regional counts in the principal components analysis presented in Figure 3. (Mandibular; Nasal; Maxillary; L – left; R – right).

**Supp Table 1.** Mk and gamma Mk analyses.

AIC weights of combined fitMk and fitGammaMk models

| model | logL | k | AIC | AIC_weight |
|---|---|---|---|---|
| One-rate | -28.598 | 1 | 59.196 | 0.204 |
| Six-rate | -25.896 | 6 | 63.794 | 0.020 |
| Ordered | -25.896 | 4 | 59.794 | 0.151 |
| Irreversible | -30.659 | 3 | 67.320 | 0.004 |
| One-rate-gamma | -26.842 | 2 | 57.684 | 0.435 |
| Six-rate-gamma | -25.041 | 7 | 64.084 | 0.018 |
| Ordered-gamma | -25.041 | 5 | 60.084 | 0.131 |
| Irreversible-gamma | -27.311 | 4 | 62.622 | 0.037 |

**Supp Table 2.** Stochastic Mapping Analyses

Stochastic Mapping Analyses

Each of the four analyses was run for 10,000 generations and sampled every 10 generations, yielding 1000 samples.

OneRate Gamma Mk Model

| State_Change | Min | Median | Mean | Max | 95% HPD |
|---|---|---|---|---|---|
| Biphasic->Subterranean | 0 | 6 | 6.948 | 33 | 0-16 |
| Biphasic->Surface | 0 | 6 | 7.525 | 31 | 0-17 |
| Subterranean->Biphasic | 0 | 5 | 6.337 | 30 | 0-15 |
| Subterranean->Surface | 0 | 6 | 7.390 | 28 | 0-17 |
| Surface->Biphasic | 1 | 7 | 7.331 | 29 | 1-16 |
| Surface->Subterranean | 2 | 9 | 9.484 | 28 | 4-18 |

SixRate Gamma Mk Model

| State_Change | Min | Median | Mean | Max | 95% HPD |
|---|---|---|---|---|---|
| Biphasic->Subterranean | 0 | 1 | 1.862 | 10 | 0-5 |
| Biphasic->Surface | 0 | 2 | 2.895 | 16 | 0-8 |
| Subterranean->Biphasic | 0 | 1 | 1.503 | 10 | 0-6 |
| Subterranean->Surface | 0 | 3 | 3.743 | 17 | 0-10 |
| Surface->Biphasic | 0 | 2 | 2.152 | 11 | 0-5 |
| Surface->Subterranean | 1 | 7 | 7.313 | 18 | 312 |

Ordered Gamma Mk Model

| State_Change | Min | Median | Mean | Max | 95% HPD |
| --- | --- | --- | --- | --- | --- |
| Biphasic->Subterranean | 0 | 0 | 0.000 | 0 | 0-0 |
| Biphasic->Surface | 2 | 3 | 3.609 | 16 | 2-7 |
| Subterranean->Biphasic | 0 | 0 | 0.000 | 0 | 0-0 |
| Subterranean->Surface | 0 | 3 | 3.386 | 19 | 0-10 |
| Surface->Biphasic | 0 | 2 | 2.281 | 15 | 1-6 |
| Surface->Subterranean | 4 | 7 | 7.844 | 19 | 6-12 |

Irreversible Gamma Mk Model

| State_Change | Min | Median | Mean | Max | 95% HPD |
| --- | --- | --- | --- | --- | --- |
| Biphasic->Subterranean | 0 | 0 | 0.000 | 0 | 0-0 |
| Biphasic->Surface | 6 | 6 | 6.301 | 9 | 6-7 |
| Subterranean->Biphasic | 0 | 0 | 0.000 | 0 | 0-0 |
| Subterranean->Surface | 0 | 1 | 1.143 | 9 | 0-4 |
| Surface->Biphasic | 0 | 0 | 0.000 | 0 | 0-0 |
| Surface->Subterranean | 6 | 7 | 7.212 | 13 | 6-9 |

**Supp Table 3.** Phylogenetic signal results

Results for Retina Total Volume
Lambda = 0.722
P(lambda = 0) = 0.0368
P(lambda = 1) = < 0.0001

Results for Lens Total Volume
Lambda = 0.952
P(lambda = 0) = < 0.0001
P(lambda = 1) = 0.015726

Results for Eye Total Volume
Lambda = 0.861
P(lambda = 0) = 0.000425
P(lambda = 1) = 0.000117

Results for PCI
Lambda = 0.853
P(lambda = 0) = 0.00125
P(lambda = 1) = 0.000125

Results for PCII
Lambda = 0
P(lambda = 0) = 1
P(lambda = 1) = < 0.0001

Results for PCIII
Lambda = 0.503
P(lambda = 0) = 0.194
P(lambda = 1) = < 0.0001

Results for PPCI
Lambda = 0.892
P(lambda = 0) = 0.000131
P(lambda = 1) = 0.000452

Results for PPCII
Lambda = 0
P(lambda = 0) = 1.000
P(lambda = 1) = < 0.0001

Results for PPCIII
Lambda = 0.986
P(lambda = 0) = < 0.0001
P(lambda = 1) = 0.853

**Supp Table 4.** Standard Principal Components AnalysisComponent Results

Eigenvalues

| Variables | PCI | PCII | PCIII |
|---|---|---|---|
| Eigenvalues | 1.924 | 0.937 | 0.139 |
| Variance Explained | 0.641 | 0.312 | 0.046 |

Loadings

| Variables | PCI | PCII | PCIII |
|---|---|---|---|
| Retina Total Volume | 0.685 | -0.173 | -0.708 |
| Lens Total Volume | 0.683 | -0.184 | 0.706 |
| SGL | 0.253 | 0.967 | 0.008 |

Phylogenetic Principal Components Analysis

Eigenvalues

| Variables | PPCI | PPCII | PPCIII |
|---|---|---|---|
| Eigenvalues | 1.908 | 0.966 | 0.126 |
| Variance Explained | 0.636 | 0.322 | 0.042 |

Loadings

| Variables | PPCI | PPCII | PPCIII |
|---|---|---|---|
| Retina Total Volume | 0.951 | -0.185 | 0.249 |
| Lens Total Volume | 0.963 | -0.093 | -0.252 |
| SGL | 0.276 | 0.961 | 0.023 |

**Supp Table 5.** Standard and Phylogenetic ANOVA/ANCOVA: Eye ~ LifeHistory. Covariate: SGL. df1 and df2 are degrees of freedom Results for F-test.Eye

Table: Standard and Phylogenetic ANOVA/ANCOVA: Eye ~ LifeHistory. Covariate: SGL.
df1 and df2 are degrees of freedom for F-test.

| Modnames | K | AIC | Delta_AIC | ModelLik | AICWt | LL | Cum.Wt | F | df1 | df2 | p |
|---|---|---|---|---|---|---|---|---|---|---|---|
| GLS_anova | 4 | 66.6572 | 0.0000 | 1.0000 | 0.5535 | -29.3286 | 0.5535 | 22.8205 | 2 | 23 | 0 |
| GLS_ancova | 5 | 67.0911 | 0.4339 | 0.8050 | 0.4456 | -28.5455 | 0.9991 | 21.8747 | 2 | 22 | 0 |
| PGLS_anova | 4 | 80.1624 | 13.5052 | 0.0012 | 0.0006 | -36.0812 | 0.9998 | 22.3119 | 2 | 23 | 0 |
| PGLS_ancova | 5 | 82.1003 | 15.4432 | 0.0004 | 0.0002 | -36.0502 | 1.0000 | 22.2698 | 2 | 22 | 0 |

**Post-hoc Comparisons: GLS_anova**

| contrast | null.value | estimate | std.error | statistic | adj.p.value |
|---|---|---|---|---|---|
| Subterranean - Biphasic | 0 | -2.9575 | 0.4433 | -6.6710 | 0.0000 |
| Surface - Biphasic | 0 | -1.7924 | 0.4231 | -4.2364 | 0.0001 |
| Surface - Subterranean | 0 | 1.1652 | 0.3505 | 3.3244 | 0.0025 |

**Post-hoc Comparisons: GLS_ancova**

| contrast | null.value | estimate | std.error | statistic | adj.p.value |
|---|---|---|---|---|---|
| Subterranean - Biphasic | 0 | -2.9305 | 0.4405 | -6.6531 | 0.0000 |
| Surface - Biphasic | 0 | -1.7325 | 0.4229 | -4.0968 | 0.0001 |
| Surface - Subterranean | 0 | 1.1980 | 0.3489 | 3.4339 | 0.0017 |

**Post-hoc Comparisons: PGLS_anova**

| contrast | null.value | estimate | std.error | statistic | adj.p.value |
|---|---|---|---|---|---|

| | | | | | |
|---|---|---|---|---|---|
| Subterranean - Biphasic | 0 | -1.8986 | 0.3037 | -6.2508 | 0.0000 |
| Surface - Biphasic | 0 | -0.7350 | 0.2454 | -2.9949 | 0.0073 |
| Surface - Subterranean | 0 | 1.1636 | 0.2016 | 5.7721 | 0.0000 |

**Post-hoc Comparisons: PGLS_ancova**

| contrast | null.value | estimate | std.error | statistic | adj.p.value |
|---|---|---|---|---|---|
| Subterranean - Biphasic | 0 | -1.8741 | 0.3280 | -5.7134 | 0.000 |
| Surface - Biphasic | 0 | -0.6899 | 0.3186 | -2.1656 | 0.075 |
| Surface - Subterranean | 0 | 1.1842 | 0.2247 | 5.2707 | |

**Supp Table 6.** Standard and Phylogenetic ANOVA/ANCOVA: Retina ~ LifeHistory. Covariate: SGL. df1 and df2 are degrees of freedom for F-test.

| Modnames | K | AIC | Delta_AIC | ModelLik | AICWt | LL | Cum.Wt | F | df1 | df2 | p |
|---|---|---|---|---|---|---|---|---|---|---|---|
| GLS_ancova | 5 | 18.6996 | 0.0000 | 1.0000 | 0.5814 | -4.3498 | 0.5814 | 42.7631 | 2 | 23 | 0 |
| GLS_anova | 4 | 19.3565 | 0.6568 | 0.7201 | 0.4186 | -5.6782 | 1.0000 | 41.0231 | 2 | 22 | 0 |
| PGLS_anova | 4 | 40.8763 | 22.1766 | 0.0000 | 0.0000 | -16.4381 | 1.0000 | 39.6187 | 2 | 23 | 0 |
| PGLS_ancova | 5 | 42.8165 | 24.1169 | 0.0000 | 0.0000 | -16.4083 | 1.0000 | 41.6601 | 2 | 22 | 0 |

Post-hoc Comparisons: PGLS_anova

| contrast | null.value | estimate | std.error | statistic | adj.p.value |
|---|---|---|---|---|---|
| Subterranean - Biphasic | 0 | -1.1960 | 0.1427 | -8.3818 | 0e+00 |
| Surface - Biphasic | 0 | -0.4271 | 0.1153 | -3.7044 | 6e-04 |
| Surface - Subterranean | 0 | 0.7689 | 0.0947 | 8.1192 | 0e+00 |

Post-hoc Comparisons: PGLS_ancova

| contrast | null.value | estimate | std.error | statistic | adj.p.value |
|---|---|---|---|---|---|
| Subterranean - Biphasic | 0 | -1.1847 | 0.1541 | -7.6877 | 0.0000 |
| Surface - Biphasic | 0 | -0.4063 | 0.1497 | -2.7147 | 0.0177 |
| Surface - Subterranean | 0 | 0.7784 | 0.1056 | 7.3745 | 0.0000 |

Post-hoc Comparisons: GLS_anova

| contrast | null.value | estimate | std.error | statistic | adj.p.value |
|---|---|---|---|---|---|

```
Subterranean - Biphasic          0       -1.5411    0.1785   -
8.6324       0e+00
     Surface - Biphasic          0       -0.7172    0.1704   -
4.2097       1e-04
 Surface - Subterranean          0        0.8239    0.1411
5.8376       0e+00
```

```
Post-hoc Comparisons: GLS_ancova
               contrast      null.value   estimate  std.error
statistic    adj.p.value
Subterranean - Biphasic          0       -1.5270    0.1737   -
8.7920       0e+00
     Surface - Biphasic          0       -0.6861    0.1667   -
4.1146       1e-04
 Surface - Subterranean          0        0.8409    0.1376
6.1130       0e+00
```

**Supp Table 7**. Standard and Phylogenetic ANOVA/ANCOVA: Lens ~ LifeHistory. Covariate: SGL.
df1 and df2 are degrees of freedom for F-test

Table: Standard and Phylogenetic ANOVA/ANCOVA: Lens ~ LifeHistory. Covariate: SGL.
df1 and df2 are degrees of freedom for F-test

| Modnames | K | AIC | Delta_AIC | ModelLik | AICWt | LL | Cum.Wt | F | df1 | df2 | p |
|---|---|---|---|---|---|---|---|---|---|---|---|
| GLS_anova | 4 | 43.2345 | 0.0000 | 1.0000 | 0.6251 | -17.6172 | 0.6251 | 9.0273 | 2 | 23 | 0.0013 |
| GLS_ancova | 5 | 44.3544 | 1.1199 | 0.5712 | 0.3571 | -17.1772 | 0.9822 | 8.5781 | 2 | 22 | 0.0018 |
| PGLS_anova | 4 | 50.9933 | 7.7588 | 0.0207 | 0.0129 | -21.4967 | 0.9951 | 12.9056 | 2 | 23 | 0.0002 |
| PGLS_ancova | 5 | 52.9379 | 9.7035 | 0.0078 | 0.0049 | -21.4690 | 1.0000 | 12.4050 | 2 | 22 | 0.0002 |

Post-hoc Comparisons: PGLS_anova

| contrast | null.value | estimate | std.error | statistic | adj.p.value |
|---|---|---|---|---|---|
| Subterranean - Biphasic | 0 | -0.7026 | 0.1733 | -4.0535 | 0.0002 |
| Surface - Biphasic | 0 | -0.3079 | 0.1401 | -2.1987 | 0.0679 |
| Surface - Subterranean | 0 | 0.3947 | 0.1150 | 3.4309 | 0.0018 |

Post-hoc Comparisons: PGLS_ancova

| contrast | null.value | estimate | std.error | statistic | adj.p.value |
|---|---|---|---|---|---|
| Subterranean - Biphasic | 0 | -0.6894 | 0.1872 | -3.6825 | 0.0007 |
| Surface - Biphasic | 0 | -0.2836 | 0.1818 | -1.5598 | 0.2583 |
| Surface - Subterranean | 0 | 0.4058 | 0.1282 | 3.1646 | 0.0043 |

Post-hoc Comparisons: GLS_anova

| contrast | null.value | estimate | std.error | statistic | adj.p.value |
|---|---|---|---|---|---|
| Subterranean - Biphasic | 0 | -1.4165 | 0.2826 | -5.0129 | 0.0000 |
| Surface - Biphasic | 0 | -1.0752 | 0.2697 | -3.9873 | 0.0002 |
| Surface - Subterranean | 0 | 0.3413 | 0.2234 | 1.5277 | 0.2757 |

Post-hoc Comparisons: GLS_ancova

| contrast | null.value | estimate | std.error | statistic | adj.p.value |
|---|---|---|---|---|---|
| Subterranean - Biphasic | 0 | -1.4034 | 0.2845 | -4.9337 | 0.0000 |
| Surface - Biphasic | 0 | -1.0464 | 0.2731 | -3.8315 | 0.0004 |
| Surface - Subterranean | 0 | 0.3571 | 0.2253 | 1.5848 | 0.2501 |

**Supp Table 8.** Phylogenetic ANOVA/ANCOVA: PCI ~ LifeHistory. Covariate: SGL.
df1 and df2 are degrees of freedom for F-test

Table: Phylogenetic ANOVA/ANCOVA: PCI ~ LifeHistory. Covariate: SGL.
df1 and df2 are degrees of freedom for F-test

| Modnames | K | AIC | Delta_AIC | ModelLik | AICWt | LL | Cum.Wt | F | df1 | df2 | p |
|---|---|---|---|---|---|---|---|---|---|---|---|
| PGLS_ancova | 5 | 82.4677 | 0.0000 | 1.0000 | 0.9496 | -36.2338 | 0.9496 | 19.5682 | 2 | 23 | 0 |
| PGLS_anova | 4 | 88.3398 | 5.8721 | 0.0531 | 0.0504 | -40.1699 | 1.0000 | 22.6304 | 2 | 22 | 0 |

Post-hoc Comparisons: PGLS_anova

| contrast | null.value | estimate | std.error | statistic | adj.p.value |
|---|---|---|---|---|---|
| Subterranean - Biphasic | 0 | -2.2176 | 0.3555 | -6.2385 | 0e+00 |
| Surface - Biphasic | 0 | -1.2558 | 0.2872 | -4.3724 | 0e+00 |
| Surface - Subterranean | 0 | 0.9618 | 0.2359 | 4.0767 | 1e-04 |

Post-hoc Comparisons: PGLS_ancova

| contrast | null.value | estimate | std.error | statistic | adj.p.value |
|---|---|---|---|---|---|
| Subterranean - Biphasic | 0 | -1.9179 | 0.3303 | -5.8058 | 0.0000 |
| Surface - Biphasic | 0 | -0.7034 | 0.3208 | -2.1924 | 0.0704 |
| Surface - Subterranean | 0 | 1.2145 | 0.2263 | 5.3676 | 0.0000 |

Post-hoc Comparisons: GLS_anova

| contrast | null.value | estimate | std.error | statistic | adj.p.value |
|---|---|---|---|---|---|
| Subterranean - Biphasic | 0 | -3.0344 | 0.4999 | -6.0701 | 0.0000 |

| | | | | | |
|---|---|---|---|---|---|
| Surface - Biphasic | 0 | -1.8790 | 0.4771 | -3.9387 | 0.0002 |
| Surface - Subterranean | 0 | 1.1554 | 0.3952 | 2.9237 | 0.0095 |

Post-hoc Comparisons: GLS_ancova

| contrast | null.value | estimate | std.error | statistic | adj.p.value |
|---|---|---|---|---|---|
| Subterranean - Biphasic | 0 | -2.9701 | 0.4405 | -6.7421 | 0.0000 |
| Surface - Biphasic | 0 | -1.7367 | 0.4229 | -4.1063 | 0.0001 |
| Surface - Subterranean | 0 | 1.2334 | 0.3489 | 3.5349 | 0.0012 |

**Supp Table 9.** Standard and Phylogenetic ANOVA/ANCOVA: PCII ~ LifeHistory. Covariate: SGL.
df1 and df2 are degrees of freedom for F-test
Table: Standard and Phylogenetic ANOVA/ANCOVA: PCII ~ LifeHistory. Covariate: SGL.
df1 and df2 are degrees of freedom for F-test

| Modnames | K | AIC | Delta_AIC | ModelLik | AICWt | LL | Cum.Wt | F | df1 | df2 | p |
|---|---|---|---|---|---|---|---|---|---|---|---|
| GLS_ancova | 5 | -3.0189 | 0.0000 | 1e+00 | 0.9996 | 6.5095 | 0.9996 | 8.9539 | 2 | 23 | 0.0013 |
| PGLS_ancova | 5 | 12.6164 | 15.6353 | 4e-04 | 0.0004 | -1.3082 | 1.0000 | 23.1907 | 2 | 22 | 0.0000 |
| GLS_anova | 4 | 77.0269 | 80.0458 | 0e+00 | 0.0000 | -34.5134 | 1.0000 | 0.9401 | 2 | 23 | 0.4051 |
| PGLS_anova | 4 | 118.8191 | 121.8380 | 0e+00 | 0.0000 | -55.4095 | 1.0000 | 23.3808 | 2 | 22 | 0.0000 |

Post-hoc Comparisons: PGLS_anova

| contrast | null.value | estimate | std.error | statistic | adj.p.value |
|---|---|---|---|---|---|
| Subterranean - Biphasic | 0 | -0.5389 | 0.6388 | -0.8437 | 0.6666 |
| Surface - Biphasic | 0 | -1.7418 | 0.5161 | -3.3747 | 0.0021 |
| Surface - Subterranean | 0 | -1.2029 | 0.4240 | -2.8372 | 0.0120 |

Post-hoc Comparisons: PGLS_ancova

| contrast | null.value | estimate | std.error | statistic | adj.p.value |
|---|---|---|---|---|---|
| Subterranean - Biphasic | 0 | 0.5064 | 0.0862 | 5.8732 | 0.0000 |
| Surface - Biphasic | 0 | 0.1852 | 0.0837 | 2.2119 | 0.0672 |
| Surface - Subterranean | 0 | -0.3212 | 0.0591 | -5.4384 | 0.0000 |

Post-hoc Comparisons: GLS_anova

| contrast | null.value | estimate | std.error | statistic | adj.p.value |
| --- | --- | --- | --- | --- | --- |
| Subterranean - Biphasic | 0 | 0.6449 | 0.5412 | 1.1917 | 0.4556 |
| Surface - Biphasic | 0 | 0.1557 | 0.5165 | 0.3015 | 0.9507 |
| Surface - Subterranean | 0 | -0.4892 | 0.4278 | -1.1434 | 0.4849 |

Post-hoc Comparisons: GLS_ancova

| contrast | null.value | estimate | std.error | statistic | adj.p.value |
| --- | --- | --- | --- | --- | --- |
| Subterranean - Biphasic | 0 | 0.7787 | 0.1144 | 6.8078 | 0e+00 |
| Surface - Biphasic | 0 | 0.4517 | 0.1098 | 4.1129 | 1e-04 |
| Surface - Subterranean | 0 | -0.3270 | 0.0906 | -3.6098 | 9e-04 |

**Supp Table 10.** Phylogenetic ANOVA/ANCOVA: PCIII ~ LifeHistory. Covariate: SGL.
df1 and df2 are degrees of freedom for F-test.

Table: Phylogenetic ANOVA/ANCOVA: PCIII ~ LifeHistory. Covariate: SGL.
df1 and df2 are degrees of freedom for F-test

| Modnames | K | AIC | Delta_AIC | ModelLik | AICWt | LL | Cum.Wt | F | df1 | df2 | p |
| --- | --- | --- | --- | --- | --- | --- | --- | --- | --- | --- | --- |
| PGLS_anova | 4 | 18.5908 | 0.0000 | 1.00 | 0.7246 | -5.2954 | 0.7246 | 29.4372 | 2 | 23 | 0 |
| PGLS_ancova | 5 | 20.5258 | 1.9351 | 0.38 | 0.2754 | -5.2629 | 1.0000 | 28.1389 | 2 | 22 | 0 |

Post-hoc Comparisons: PGLS_anova

| contrast | null.value | estimate | std.error | statistic | adj.p.value |
| --- | --- | --- | --- | --- | --- |
| Subterranean - Biphasic | 0 | -0.6110 | 0.0930 | -6.5736 | 0.000 |

```
     Surface - Biphasic          0         -0.1739        0.0751
-2.3150       0.051
 Surface - Subterranean          0          0.4372        0.0617
7.0862       0.000


Post-hoc Comparisons: PGLS_ancova
               contrast      null.value    estimate
std.error     statistic     adj.p.value
Subterranean - Biphasic          0         -0.6034        0.1004
-6.0109      0.0000
     Surface - Biphasic          0         -0.1598        0.0975
-1.6387      0.2248
 Surface - Subterranean          0          0.4436        0.0688
6.4522      0.0000


Post-hoc Comparisons: GLS_anova
               contrast      null.value    estimate
std.error     statistic     adj.p.value
Subterranean - Biphasic          0         -0.2664        0.1591
-1.6742      0.2132
     Surface - Biphasic          0          0.2900        0.1518
1.9100      0.1343
 Surface - Subterranean          0          0.5564        0.1258
4.4233      0.0000


Post-hoc Comparisons: GLS_ancova
               contrast      null.value    estimate
std.error     statistic     adj.p.value
Subterranean - Biphasic          0         -0.2629        0.1623
-1.6199      0.2351
     Surface - Biphasic          0          0.2977        0.1558
1.9103      0.1343
 Surface - Subterranean          0          0.5606        0.1286
4.3607      0.0000
```

**Supp Table 11.**Log-likelihoods, degrees Results of freedom, AIC, and AIC weight for
the one-rate, two-rate, three-rate, and four-rate models, for each character
in the rate -shift analysis.

Eye

| | log(L) | df | AIC | weight |
|---|---|---|---|---|
| Rate1_Best_Rep | -44.5674 | 2 | 93.1349 | 0.0019 |
| Rate2_Best_Rep | -37.0048 | 4 | 82.0097 | 0.5127 |
| Rate3_Best_Rep | -35.3443 | 6 | 82.6886 | 0.3651 |
| Rate4_Best_Rep | -34.4556 | 8 | 84.9113 | 0.1202 |

Lens

| | log(L) | df | AIC | weight |
|---|---|---|---|---|
| Rate1_Best_Rep | -18.8273 | 2 | 41.6546 | 0.0502 |
| Rate2_Best_Rep | -15.8521 | 4 | 39.7041 | 0.1332 |
| Rate3_Best_Rep | -12.3455 | 6 | 36.6910 | 0.6009 |
| Rate4_Best_Rep | -11.3700 | 8 | 38.7401 | 0.21570 |

Retina

| | log(L) | df | AIC | weight |
|---|---|---|---|---|
| Rate1_Best_Rep | -33.7529 | 2 | 71.5058 | 8.8600e-05 |
| Rate2_Best_Rep | -22.6516 | 4 | 53.3032 | 7.9449e-01 |
| Rate3_Best_Rep | -22.3110 | 6 | 56.6220 | 1.5115e-01 |
| Rate4_Best_Rep | -21.3355 | 8 | 58.6710 | 5.4261e-02 |

SGL

| | log(L) | df | AIC | weight |
|---|---|---|---|---|
| Rate1_Best_Rep | -62.3925 | 2 | 128.7851 | 2.5283e-05 |
| Rate2_Best_Rep | -50.1337 | 4 | 108.2673 | 7.2147e-01 |
| Rate3_Best_Rep | -49.4207 | 6 | 110.8414 | 1.9919e-01 |
| Rate4_Best_Rep | -48.3415 | 8 | 112.6831 | 7.9311e-02 |

Eye_Residuals

| | log(L) | df | AIC | weight |
|---|---|---|---|---|
| Rate1_Best_Rep | -44.3690 | 2 | 92.7381 | 0.0018 |
| Rate2_Best_Rep | -36.6286 | 4 | 81.2571 | 0.5573 |
| Rate3_Best_Rep | -35.1470 | 6 | 82.2958 | 0.3316 |
| Rate4_Best_Rep | -34.2573 | 8 | 84.5147 | 0.1093 |

Lens_Residuals

| | log(L) | df | AIC | weight |
|---|---|---|---|---|
| Rate1_Best_Rep | -18.4435 | 2 | 40.8870 | 0.0632 |
| Rate2_Best_Rep | -15.3795 | 4 | 38.7591 | 0.1834 |
| Rate3_Best_Rep | -12.2791 | 6 | 36.5583 | 0.5511 |
| Rate4_Best_Rep | -11.2825 | 8 | 38.5650 | 0.2020 |

Retina_Residuals

| | log(L) | df | AIC | weight |
|---|---|---|---|---|
| Rate1_Best_Rep | -33.6460 | 2 | 71.2921 | 7.4267e-05 |
| Rate2_Best_Rep | -22.3653 | 4 | 52.7307 | 7.968e-01 |
| Rate3_Best_Rep | -22.0075 | 6 | 56.0150 | 1.54235e-01 |
| Rate4_Best_Rep | -21.1568 | 8 | 58.3136 | 4.8870e-02 |

PPC1

| | log(L) | df | AIC | weight |
|---|---|---|---|---|
| Rate1_Best_Rep | -21.9029 | 2 | 47.8059 | 0.0055 |
| Rate2_Best_Rep | -15.7282 | 4 | 39.4565 | 0.3602 |
| Rate3_Best_Rep | -13.4440 | 6 | 38.8881 | 0.4786 |
| Rate4_Best_Rep | -12.5682 | 8 | 41.1364 | 0.1555 |

PPC2

| | log(L) | df | AIC | weight |
|---|---|---|---|---|
| Rate1_Best_Rep | -13.05706 | 2 | 30.1141 | 2.9741e-05 |
| Rate2_Best_Rep | -0.97183 | 4 | 9.9436 | 7.1338e-01 |
| Rate3_Best_Rep | -0.42939 | 6 | 12.8587 | 1.6607e-01 |
| Rate4_Best_Rep | 1.24983 | 8 | 13.5003 | 1.2050e-01 |

PPC3

| | log(L) | df | AIC | weight |
|---|---|---|---|---|
| Rate1_Best_Rep | 13.4347 | 2 | -22.86956 | 0.3559361 |
| Rate2_Best_Rep | 14.2980 | 4 | -20.59608 | 0.1142068 |
| Rate3_Best_Rep | 16.9292 | 6 | -21.85849 | 0.2146947 |
| Rate4_Best_Rep | 19.3131 | 8 | -22.62623 | 0.3151625 |

**Supp Table 12**. Generalized linear model for eye volume given species and stage.

| Coefficients: | Estimate | Std. Error | t value | Pr(>\|t\|) |
|---|---|---|---|---|
| (Intercept) | 0.054161 | 0.008458 | 6.404 | 1.30e-08 *** |
| speciesE.nana | -0.028525 | 0.011961 | -2.385 | 0.01969 * |
| speciesE.pterophila_CS | 0.015655 | 0.011961 | 1.309 | 0.19470 |
| speciesE.pterophila_PC | 0.004194 | 0.011961 | 0.351 | 0.72688 |
| speciesE.rathbuni | -0.034782 | 0.011961 | -2.908 | 0.00482 ** |
| speciesE.sosorum | -0.012670 | 0.011961 | -1.059 | 0.29296 |
| stage2_month | -0.016134 | 0.011961 | -1.349 | 0.18154 |
| stage3_month | -0.021696 | 0.011961 | -1.814 | 0.07381 . |
| stage4_month | -0.023967 | 0.011961 | -2.004 | 0.04881 * |
| stage5_month | -0.023969 | 0.011961 | -2.004 | 0.04879 * |
| stageAdult | -0.028715 | 0.011961 | -2.401 | 0.01891 * |
| speciesE.nana:stage2_month | 0.078596 | 0.016916 | 4.646 | 1.46e-05 *** |
| speciesE.pterophila_CS:stage2_month | 0.045261 | 0.016916 | 2.676 | 0.00920 ** |
| speciesE.pterophila_PC:stage2_month | 0.005365 | 0.016916 | 0.317 | 0.75201 |
| speciesE.rathbuni:stage2_month | 0.011496 | 0.016916 | 0.680 | 0.49891 |
| speciesE.sosorum:stage2_month | 0.022162 | 0.016916 | 1.310 | 0.19425 |
| speciesE.nana:stage3_month | 0.099332 | 0.016916 | 5.872 | 1.18e-07 *** |
| speciesE.pterophila_CS:stage3_month | 0.046905 | 0.016916 | 2.773 | 0.00705 ** |
| speciesE.pterophila_PC:stage3_month | 0.009293 | 0.016916 | 0.549 | 0.58444 |
| speciesE.rathbuni:stage3_month | 0.011048 | 0.016916 | 0.653 | 0.51572 |

```
speciesE.sosorum:stage3_month        0.024583   0.016916   1.453
0.15044
speciesE.nana:stage4_month           0.100580   0.016916   5.946
8.71e-08 ***
speciesE.pterophila_CS:stage4_month  0.073059   0.016916   4.319
4.86e-05 ***
speciesE.pterophila_PC:stage4_month  0.002288   0.016916   0.135
0.89280
speciesE.rathbuni:stage4_month       0.013284   0.016916   0.785
0.43481
speciesE.sosorum:stage4_month        0.022255   0.016916   1.316
0.19242
speciesE.nana:stage5_month           0.110395   0.016916   6.526
7.78e-09 ***
speciesE.pterophila_CS:stage5_month  0.045353   0.016916   2.681
0.00907 **
speciesE.pterophila_PC:stage5_month  0.001621   0.016916   0.096
0.92393
speciesE.rathbuni:stage5_month       0.009104   0.016916   0.538
0.59210
speciesE.sosorum:stage5_month        0.033645   0.016916   1.989
0.05045 .
speciesE.nana:stageAdult             0.206574   0.016916  12.212
<2e-16 ***
speciesE.pterophila_CS:stageAdult    0.199341   0.016916  11.784
<2e-16 ***
speciesE.pterophila_PC:stageAdult   -0.001846   0.016379  -0.113
0.91055
speciesE.rathbuni:stageAdult         0.014670   0.016916   0.867
0.38866
speciesE.sosorum:stageAdult          0.112368   0.016916   6.643
4.75e-09 ***
```

**Supp. Table 13a.** 1-month post oviposition linear model results.

| | Estimate | Std. Error | t value | Pr(>\|t\|) | |
|---|---|---|---|---|---|
| (Intercept) | 0.054161 | 0.054161 | 12.606 | 2.79e-08 | *** |
| speciesE.nana | -0.028525 | 0.006076 | -4.695 | 0.000519 | *** |
| speciesE.pterophila_CS | 0.015655 | 0.006076 | 2.577 | 0.024250 | * |
| speciesE.pterophila_PC | 0.004194 | 0.006076 | 0.690 | 0.503170 | |
| speciesE.rathbuni | -0.034782 | 0.006076 | -5.725 | 9.54e-05 | *** |
| speciesE.sosorum | -0.012670 | 0.006076 | -2.085 | 0.059061 | |

**Supp. Table 13b.** 1-month post oviposition ANOVA results.

| | Df | Sum Sq | Mean Sq | F value | Pr(>F) | |
|---|---|---|---|---|---|---|
| species | 5 | 0.005765 | 0.0011530 | 20.82 | 1.56e-05 | *** |
| Residuals | 12 | 0.000664 | 0.0000554 | | | |

**Supp. Table 13c.** 1- month post oviposition Tukey's results with estimated difference in means (diff) and upper (upr) and lower (lwr) bounds of the confidence interval.

| | diff | lwr | upr | p adj |
|---|---|---|---|---|
| E.nana-E.latitans_HCC | -0.0285 | -0.048933412 | -0.008116508 | 0.0053186 |
| E.pterophila_CS-E.latitans_HCC | 0.015 | -0.004753472 | 0.036063432 | 0.1768363 |
| E.pterophila_PC-E.latitans_HCC | 0.004 | -0.016214572 | 0.024602332 | 0.9796568 |
| E.rathbuni-E.latitans_HCC | -0.034 | -0.055190552 | -0.014373648 | 0.0010344 |

| | | | | |
|---|---|---|---|---|
| E.sosorum-E.latitans_HCC | -0.012 | -0.033078785 | 0.007738118 | 0.3549104 |
| E.pterophila_CS-E.nana | 0.04417994 | 0.023771488 | 0.064588392 | 0.0001119 |
| E.pterophila_PC-E.nana | 0.03271884 | 0.012310388 | 0.053127292 | 0.0017520 |
| E.rathbuni-E.nana | -0.00625714 | -0.026665592 | 0.014151312 | 0.8989568 |
| E.sosorum-E.nana | 0.01585463 | -0.004553825 | 0.036263078 | 0.1681852 |
| E.pterophila_PC-E.pterophila_CS | -0.01146110 | -0.031869552 | 0.008947352 | 0.4534947 |
| E.rathbuni-E.pterophila_CS | -0.05043708 | -0.070845532 | -0.030028628 | 0.0000297 |
| E.sosorum-E.pterophila_CS | -0.02832531 | -0.048733765 | -0.007916862 | 0.0056140 |
| E.rathbuni-E.pterophila_PC | -0.03897598 | -0.059384432 | -0.018567528 | 0.0003699 |
| E.sosorum-E.pterophila_PC | -0.01686421 | -0.037272665 | 0.003544238 | 0.1298459 |
| E.sosorum-E.rathbuni | 0.02211177 | 0.001703315 | 0.042520218 | 0.0311923 |

**Supp. Table 13d.** 1-month post oviposition Tukey's Grouping.

| | vol. | groups |
|---|---|---|
| E.pterophila_CS | 0.06981598 | a |
| E.pterophila_PC | 0.05835488 | ab |
| E.latitans_HCC | 0.05416100 | ab |
| E.sosorum | 0.04149067 | bc |
| E.nana | 0.02563604 | cd |
| E.rathbuni | 0.01937890 | d |

**Supp. Table 14a.** 2-month post oviposition linear model results.

| | Estimate | Std. Error | t-value | Pr(>\|t\|) | |
|---|---|---|---|---|---|
| (Intercept) | 0.038027 | 0.004880 | 7.792 | 4.92e-06 | *** |
| speciesE.nana | 0.050071 | 0.006902 | 7.255 | 1.01e-05 | *** |
| speciesE.pterophila_CS | 0.060916 | 0.006902 | 8.826 | 1.36e-06 | *** |
| speciesE.pterophila_PC | 0.009559 | 0.006902 | 1.385 | 0.19125 | |
| speciesE.rathbuni | -0.023286 | 0.006902 | -3.374 | 0.00553 | ** |
| speciesE.sosorum | 0.009492 | 0.006902 | 1.375 | 0.19418 | |

**Supp. Table 14b.** 2-month post oviposition ANOVA.

| | Df | Sum Sq | Mean Sq | F value | Pr(>F) | |
|---|---|---|---|---|---|---|
| species | 5 | 0.015127 | 0.0030253 | 42.34 | 3.26e-07 | *** |
| Residuals | 12 | 0.000857 | 0.0000715 | | | |

**Supp. Table 14c.** 2-month post oviposition Tukey's results with estimated difference in means (diff), upper (upr) and lower (lwr) bounds of the confidence interval, and p adjusted p-value.

| | diff | lwr | upr |
|---|---|---|---|
| E.nana-E.latitans_HCC | 5.007063e-02 | 0.026887817 | 0.0732534493 |
| E.pterophila_CS-E.latitans_HCC | 6.091603e-02 | 0.037733217 | 0.0840988493 |
| E.pterophila_PC-E.latitans_HCC | 9.559333e-03 | -0.013623483 | 0.0327421493 |
| E.rathbuni-E.latitans_HCC | -2.328615e-02 | -0.046468963 | -0.0001033307 |
| E.sosorum-E.latitans_HCC | 9.492000e-03 | -0.013690816 | 0.0326748160 |
| E.pterophila_CS-E.nana | 1.084540e-02 | -0.012337416 | 0.0340282160 |
| E.pterophila_PC-E.nana | -4.051130e-02 | -0.063694116 | -0.0173284840 |
| E.rathbuni-E.nana | -7.335678e-02 | -0.096539596 | -0.0501739640 |
| E.sosorum-E.nana | -4.057863e-02 | -0.063761449 | -0.0173958173 |
| E.pterophila_PC-E.pterophila_CS | -5.135670e-02 | -0.074539516 | -0.0281738840 |
| E.rathbuni-E.pterophila_CS | -8.420218e-02 | -0.107384996 | -0.0610193640 |
| E.sosorum-E.pterophila_CS | -5.142403e-02 | -0.074606849 | -0.0282412173 |
| E.rathbuni-E.pterophila_PC | -3.284548e-02 | -0.056028296 | -0.0096626640 |
| E.sosorum-E.pterophila_PC | -6.733333e-05 | -0.023250149 | 0.0231154827 |
| E.sosorum-E.rathbuni | 3.277815e-02 | 0.009595331 | 0.0559609627 |

p adj

| | |
|---|---|
| E.nana-E.latitans_HCC | 0.0001144 |
| E.pterophila_CS-E.latitans_HCC | 0.0000158 |
| E.pterophila_PC-E.latitans_HCC | 0.7348996 |
| E.rathbuni-E.latitans_HCC | 0.0487616 |
| E.sosorum-E.latitans_HCC | 0.7401905 |
| E.pterophila_CS-E.nana | 0.6299014 |
| E.pterophila_PC-E.nana | 0.0008294 |
| E.rathbuni-E.nana | 0.0000022 |
| E.sosorum-E.nana | 0.0008173 |
| E.pterophila_PC-E.pterophila_CS | 0.0000892 |
| E.rathbuni-E.pterophila_CS | 0.0000005 |
| E.sosorum-E.pterophila_CS | 0.0000880 |
| E.rathbuni-E.pterophila_PC | 0.0047874 |
| E.sosorum-E.pterophila_PC | 1.0000000 |
| E.sosorum-E.rathbuni | 0.0048645 |

**Supp. Table 14d.** 2-month post oviposition Tukey's Grouping.

| | vol. | groups |
|---|---|---|
| E.pterophila_CS | 0.09894270 | a |
| E.nana | 0.08809730 | a |
| E.pterophila_PC | 0.04758600 | b |
| E.sosorum | 0.04751867 | b |
| E.latitans_HCC | 0.03802667 | b |
| E.rathbuni | 0.01474052 | c |

**Supp. Table 15a.** 3-month post oviposition linear model results.

| | Estimate | Std. Error | t value | Pr(>\|t\|) | |
|---|---|---|---|---|---|
| (Intercept) | 0.032465 | 0.003438 | 9.444 | 6.62e-07 | *** |
| speciesE.nana | 0.070807 | 0.004862 | 14.564 | 5.44e-09 | *** |
| speciesE.pterophila_CS | 0.062560 | 0.004862 | 12.868 | 2.21e-08 | *** |
| speciesE.pterophila_PC | 0.013487 | 0.004862 | 2.774 | 0.016834 | * |
| speciesE.rathbuni | -0.023734 | 0.004862 | -4.882 | 0.000377 | *** |
| speciesE.sosorum | 0.011913 | 0.004862 | 2.450 | 0.030574 | * |

**Supp. Table 15b.** 3-month post oviposition ANOVA.

| | Df | Sum Sq | Mean Sq | F value | Pr(>F) | |
|---|---|---|---|---|---|---|
| species | 5 | 0.020327 | 0.004065 | 114.7 | 1.06e-09 | *** |
| Residuals | 12 | 0.000425 | 0.000035 | | | |

**Supp. Table 15c.** 3- month post oviposition Tukey's results with estimated difference in means (diff), upper (upr) and lower (lwr) bounds of the confidence interval, and p adjusted p-value.

| | diff | lwr | upr | p adj |
|---|---|---|---|---|
| E.nana-E.latitans_HCC | 0.070806820 | 0.054476980 | 0.08713666 | 0.0000001 |
| E.pterophila_CS-E.latitans_HCC | 0.062559980 | 0.046230140 | 0.07888982 | 0.0000003 |
| E.pterophila_PC-E.latitans_HCC | 0.013486640 | -0.002843200 | 0.02981648 | 0.1301532 |
| E.rathbuni-E.latitans_HCC | -0.023733900 | -0.040063740 | -0.00740406 | 0.0039172 |
| E.sosorum-E.latitans_HCC | 0.011912667 | -0.004417173 | 0.02824251 | 0.2136540 |

| | | | | |
|---|---|---|---|---|
| E.pterophila_CS-E.nana | -0.008246840 | -0.024576680 | 0.00808300 | 0.5582512 |
| E.pterophila_PC-E.nana | -0.057320180 | -0.073650020 | -0.04099034 | 0.0000007 |
| E.rathbuni-E.nana | -0.094540720 | -0.110870560 | -0.07821088 | 0.0000000 |
| E.sosorum-E.nana | -0.058894153 | -0.075223993 | -0.04256431 | 0.0000005 |
| E.pterophila_PC-E.pterophila_CS | -0.049073340 | -0.065403180 | -0.03274350 | 0.0000038 |
| E.rathbuni-E.pterophila_CS | -0.086293880 | -0.102623720 | -0.06996404 | 0.0000000 |
| E.sosorum-E.pterophila_CS | -0.050647313 | -0.066977153 | -0.03431747 | 0.0000027 |
| E.rathbuni-E.pterophila_PC | -0.037220540 | -0.053550380 | -0.02089070 | 0.0000672 |
| E.sosorum-E.pterophila_PC | -0.001573973 | -0.017903813 | 0.01475587 | 0.9993912 |
| E.sosorum-E.rathbuni | 0.035646567 | 0.019316727 | 0.05197641 | 0.0001031 |

**Supp. Table 15d.** 3-month post oviposition Tukey’s Grouping.

| | vol. | groups |
|---|---|---|
| E.nana | 0.10327182 | a |
| E.pterophila_CS | 0.09502498 | a |
| E.pterophila_PC | 0.04595164 | b |
| E.sosorum | 0.04437767 | b |
| E.latitans_HCC | 0.03246500 | b |
| E.rathbuni | 0.00873110 | c |

**Supp. Table 16a.** 4-month post oviposition linear model results.

| | Estimate | Std. Error | t value | Pr(>\|t\|) |
|---|---|---|---|---|
| (Intercept) | 0.030194 | 0.006253 | 4.829 | 0.000413 *** |
| speciesE.nana | 0.072055 | 0.008843 | 8.149 | 3.11e-06 *** |
| speciesE.pterophila_CS | 0.088713 | 0.008843 | 10.032 | 3.46e-07 *** |
| speciesE.pterophila_PC | 0.006482 | 0.008843 | 0.733 | 0.477645 |
| speciesE.rathbuni | -0.021498 | 0.008843 | -2.431 | 0.031665 * |
| speciesE.sosorum | 0.009584 | 0.008843 | 1.084 | 0.299718 |

**Supp. Table 16b.** 4-month post oviposition ANOVA.

| | Df | Sum Sq | Mean Sq | F value | Pr(>F) |
|---|---|---|---|---|---|
| species | 5 | 0.028909 | 0.005782 | 49.3 | 1.38e-07 *** |
| Residuals | 12 | 0.001407 | 0.000117 | | |

**Supp. Table 16c.** 4- month post oviposition Tukey's results with estimated difference in means (diff), upper (upr) and lower (lwr) bounds of the confidence interval, and p adjusted p-value.

| | diff | lwr | upr | p adj |
|---|---|---|---|---|
| E.nana-E.latitans_HCC | 0.072054607 | 0.042352754 | 0.101756460 | 0.0000359 |
| E.pterophila_CS-E.latitans_HCC | 0.088713487 | 0.059011634 | 0.118415340 | 0.0000041 |
| E.pterophila_PC-E.latitans_HCC | 0.006481507 | -0.023220346 | 0.036183360 | 0.9736968 |

| | | | | |
|---|---|---|---|---|
| E.rathbuni-E.latitans_HCC | -0.021498053 | -0.051199906 | 0.008203800 | 0.2197638 |
| E.sosorum-E.latitans_HCC | 0.009584333 | -0.020117520 | 0.039286186 | 0.8787145 |
| E.pterophila_CS-E.nana | 0.016658880 | -0.013042973 | 0.046360733 | 0.4547658 |
| E.pterophila_PC-E.nana | -0.065573100 | -0.095274953 | -0.035871247 | 0.0000922 |
| E.rathbuni-E.nana | -0.093552660 | -0.123254513 | -0.063850807 | 0.0000023 |
| E.sosorum-E.nana | -0.062470273 | -0.092172126 | -0.032768420 | 0.0001481 |
| E.pterophila_PC-E.pterophila_CS | -0.082231980 | -0.111933833 | -0.052530127 | 0.0000091 |
| E.rathbuni-E.pterophila_CS | -0.110211540 | -0.139913393 | -0.080509687 | 0.0000004 |
| E.sosorum-E.pterophila_CS | -0.079129153 | -0.108831006 | -0.049427300 | 0.0000137 |
| E.rathbuni-E.pterophila_PC | -0.027979560 | -0.057681413 | 0.001722293 | 0.0691558 |
| E.sosorum-E.pterophila_PC | 0.003102827 | -0.026599026 | 0.032804680 | 0.9991023 |
| E.sosorum-E.rathbuni | 0.031082387 | 0.001380534 | 0.060784240 | 0.0384643 |

**Supp. Table 16d.** 4-month post oviposition Tukey's Grouping.

| | vol. | groups |
|---|---|---|
| E.pterophila_CS | 0.11890782 | a |
| E.nana | 0.10224894 | a |
| E.sosorum | 0.03977867 | b |
| E.pterophila_PC | 0.03667584 | bc |
| E.latitans_HCC | 0.03019433 | bc |
| E.rathbuni | 0.00869628 | c |

**Supp. Table 17a.** 5-month post oviposition linear model results.

| | Estimate | Std. Error | t value | Pr(>\|t\|) |
|---|---|---|---|---|
| (Intercept) | 0.030192 | 0.004689 | 6.439 | 3.21e-05 *** |
| speciesE.nana | 0.081870 | 0.006631 | 12.347 | 3.52e-08 *** |
| speciesE.pterophila_CS | 0.061008 | 0.006631 | 9.201 | 8.74e-07 *** |
| speciesE.pterophila_PC | 0.005815 | 0.006631 | 0.877 | 0.39773 |
| speciesE.rathbuni | -0.025679 | 0.006631 | -3.873 | 0.00222 ** |
| speciesE.sosorum | 0.020975 | 0.006631 | 3.163 | 0.00817 ** |

**Supp. Table 17b.** 5-month post oviposition ANOVA.

| | Df | Sum Sq | Mean Sq | F value | Pr(>F) |
|---|---|---|---|---|---|
| species | 5 | 0.024307 | 0.004861 | 73.72 | 1.38e-08 *** |
| Residuals | 12 | 0.000791 | 0.000066 | | |

**Supp. Table 17c.** 5-month post oviposition Tukey's results with estimated difference in means (diff), upper (upr) and lower (lwr) bounds of the confidence interval, and p adjusted p-value.

| | diff | lwr | upr | p adj |
|---|---|---|---|---|
| E.nana-E.latitans_HCC | 0.081869673 | 0.059598156 | 0.104141190 | 0.0000004 |
| E.pterophila_CS-E.latitans_HCC | 0.061008053 | 0.038736536 | 0.083279570 | 0.0000102 |
| E.pterophila_PC-E.latitans_HCC | 0.005814653 | -0.016456864 | 0.028086170 | 0.9451098 |

| | | | | |
|---|---|---|---|---|
| E.rathbuni-E.latitans_HCC | -0.025678527 | -0.047950044 | -0.003407010 | 0.0210207 |
| E.sosorum-E.latitans_HCC | 0.020975000 | -0.001296517 | 0.043246517 | 0.0692435 |
| E.pterophila_CS-E.nana | -0.020861620 | -0.043133137 | 0.001409897 | 0.0712286 |
| E.pterophila_PC-E.nana | -0.076055020 | -0.098326537 | -0.053783503 | 0.0000009 |
| E.rathbuni-E.nana | -0.107548200 | -0.129819717 | -0.085276683 | 0.0000000 |
| E.sosorum-E.nana | -0.060894673 | -0.083166190 | -0.038623156 | 0.0000104 |
| E.pterophila_PC-E.pterophila_CS | -0.055193400 | -0.077464917 | -0.032921883 | 0.0000289 |
| E.rathbuni-E.pterophila_CS | -0.086686580 | -0.108958097 | -0.064415063 | 0.0000002 |
| E.sosorum-E.pterophila_CS | -0.040033053 | -0.062304570 | -0.017761536 | 0.0006442 |
| E.rathbuni-E.pterophila_PC | -0.031493180 | -0.053764697 | -0.009221663 | 0.0048603 |
| E.sosorum-E.pterophila_PC | 0.015160347 | -0.007111170 | 0.037431864 | 0.2705121 |
| E.sosorum-E.rathbuni | 0.046653527 | 0.024382010 | 0.068925044 | 0.0001540 |

**Supp. Table 17d.** 5-month post oviposition Tukey's Grouping.

| | vol. | groups |
|---|---|---|
| E.nana | 0.11206134 | a |
| E.pterophila_CS | 0.09119972 | a |
| E.sosorum | 0.05116667 | b |
| E.pterophila_PC | 0.03600632 | b |
| E.latitans_HCC | 0.03019167 | b |
| E.rathbuni | 0.00451314 | c |

**Supp. Table 18a.** Adult linear model results.

| | Estimate | Std. Error | t value | Pr(>\|t\|) |
|---|---|---|---|---|
| (Intercept) | 0.025446 | 0.017187 | 1.481 | 0.16255 |
| speciesE.nana | 0.178049 | 0.024306 | 7.325 | 5.79e-06 *** |
| speciesE.pterophila_CS | 0.214996 | 0.024306 | 8.845 | 7.32e-07 *** |
| speciesE.pterophila_PC | 0.002347 | 0.022736 | 0.103 | 0.91934 |
| speciesE.rathbuni | -0.020112 | 0.024306 | -0.827 | 0.42290 |
| speciesE.sosorum | 0.099698 | 0.024306 | 4.102 | 0.00125 ** |

**Supp. Table 18b.** Adult ANOVA.

| | Df | Sum Sq | Mean Sq | F value | Pr(>F) |
|---|---|---|---|---|---|
| species | 5 | 0.15761 | 0.031522 | 35.57 | 3.86e-07 *** |
| Residuals | 13 | 0.01152 | 0.000886 | | |

**Supp. Table 18c.** Adult Tukey's results with estimated difference in means (diff), upper (upr) and lower (lwr) bounds of the confidence interval, and p adjusted p-value.

| | diff | lwr | upr | p adj |
|---|---|---|---|---|
| E.nana-E.latitans_HCC | 0.178049136 | 0.09744825 | 0.258650028 | 0.0000679 |
| E.pterophila_CS-E.latitans_HCC | 0.214996415 | 0.13439552 | 0.295597306 | 0.0000088 |
| E.pterophila_PC-E.latitans_HCC | 0.002347499 | -0.07304773 | 0.077742729 | 0.9999979 |

| | | | | |
|---|---|---|---|---|
| E.rathbuni-E.latitans_HCC | -0.020112202 | -0.10071309 | 0.060488689 | 0.9569322 |
| E.sosorum-E.latitans_HCC | 0.099697834 | 0.01909694 | 0.180298725 | 0.0125240 |
| E.pterophila_CS-E.nana | 0.036947279 | -0.04365361 | 0.117548170 | 0.6586908 |
| E.pterophila_PC-E.nana | -0.175701637 | -0.25109687 | -0.100306407 | 0.0000385 |
| E.rathbuni-E.nana | -0.198161339 | -0.27876223 | -0.117560447 | 0.0000216 |
| E.sosorum-E.nana | -0.078351303 | -0.15895219 | 0.002249589 | 0.0586988 |
| E.pterophila_PC-E.pterophila_CS | -0.212648916 | -0.28804415 | -0.137253686 | 0.0000047 |
| E.rathbuni-E.pterophila_CS | -0.235108617 | -0.31570951 | -0.154507726 | 0.0000032 |
| E.sosorum-E.pterophila_CS | -0.115298581 | -0.19589947 | -0.034697690 | 0.0040713 |
| E.rathbuni-E.pterophila_PC | -0.022459701 | -0.09785493 | 0.052935528 | 0.9138823 |
| E.sosorum-E.pterophila_PC | 0.097350335 | 0.02195510 | 0.172745564 | 0.0091183 |
| E.sosorum-E.rathbuni | 0.119810036 | 0.03920914 | 0.200410927 | 0.0029583 |

**Supp. Table 18d.** Adult Tukey's Grouping.

| | vol. | groups |
|---|---|---|
| E.pterophila_CS | 0.240442185 | a |
| E.nana | 0.203494907 | ab |
| E.sosorum | 0.125143604 | b |
| E.pterophila_PC | 0.027793269 | c |
| E.latitans_HCC | 0.025445770 | c |
| E.rathbuni | 0.005333568 | c |

**Supp Table 19.** List of specimens and corresponding Genbank accession numbers.

| Species | cytb | RAG1 |
|---|---|---|
| *Eurycea wallacei* | KY073012 | KF562693 |
| *Eurycea aquatica* | KF562543 | KF562645 |
| *Eurycea cirrigera* | JQ920622 | KF562650 |
| *Eurycea hillisi* | KY502648 | -- |
| *Eurycea paludicola* | AY528401 | KF562662 |
| *Eurycea* sp. 5 | TNHC 114070 | -- |
| *Eurycea tonkawae* | AY014842 | AY691709 |
| *Eurycea chisholmensis* | AY014841 | KF562647 |
| *Eurycea naufragia* | AY014843 | KF562657 |
| *Eurycea* sp. 3 metamorphic | KC355884 | -- |
| *Eurycea* sp. 3 paedomorphic | TNHC 66522 | -- |
| *Eurycea* sp. 2 | KC355910 | -- |
| *Eurycea troglodytes* subterranean | KC355939 | KF562671 |
| *Eurycea troglodytes* surface | AY014853 | KF562672 |
| *Eurycea* sp. 4 | SMARC MV | KY073109 |
| *Eurycea rathbuni* | AY014844 | AY691708 |
| *Eurycea waterlooensis* | AY014856 | KF562679 |
| *Eurycea sosorum* | KC355865 | KF562664 |
| *Eurycea nana* | AY014846 | KF562656 |
| *Eurycea* sp. 1 | AY014847 | -- |
| *Eurycea* sp. 1 Cave | AGG 2061 | -- |
| *Eurycea neotenes* | AY528400 | AY650122 |
| *Eurycea pterophila* | AY014851 | KF562658 |

| | | |
|---|---|---|
| *Eurycea pterophila* Grapevine Cave | KC355913 | -- |
| *Eurycea latitans* Honey Creek Cave | KC355918 | KF562652 |
| *Eurycea latitans* Honey Creek SNA Spg | KC355923 | -- |
